# Changes in dysbiosis and gene expression in the gut of wharf roach (*Ligia* spp.) fed with expanded polystyrene

**DOI:** 10.64898/2026.03.31.715705

**Authors:** Seokhyun Lee, Hirokuni Miyamoto, Yuki Takai, Wataru Suda, Hiroshi Ohno, Yohei Simasaki, Yuji Oshima

## Abstract

The East Asian region, known for its high levels of human and fishery activities, experiences serious plastic pollution in the marine environment, especially in seawater and along coastlines. Wharf roaches (*Ligia* spp.) collected from the coast of western Japan frequently ingest expanded polystyrene (EPS), which is then excreted as microplastic through their feces. However, the impact of EPS exposure and ingestion on the gut microbiome of wharf roaches remains unclear. Thus, this study aimed to investigate the effects of EPS ingestion on the gut microbiota of wharf roaches by examining their gut microbiota and gene expression. The expression levels of more than 400 genes, including those associated with xenobiotic metabolism, and the abundance of gut microbial community were altered. Microbial analysis revealed that at least five archaeal types, two to four bacterial types, three to seven eukaryotic types, and three viral types were involved in a correlation network composed of strong associations. Among them, *Haloquadratum*, *Halalkalicoccus*, and *Methanospirillum* (archaea); *Volvox* (eukaryote); and *Varicellovirus* and T4-like viruses showed significantly increased abundance. Furthermore, covariance structure analysis indicated that the viruses and methanogens played key causal roles as characteristic factors related to EPS administration. In conclusion, EPS disrupts the intestinal environment of wharf roaches and serves as a potential material for viral activation and methane production. Building on our previous field study that identified wharf roaches as potential indicators of coastal EPS pollution, this study provides novel insights into the ecological impacts of EPS ingestion and consequences of plastic pollution.

## Introduction

Plastic pollution in marine ecosystems is a serious environmental concern worldwide. Since the 1950s, plastic production has significantly increased. Plastic waste has become ubiquitous, manifesting in coastal areas, the deep sea, the atmosphere, and tropical and Arctic regions (MacLeod et al., 2021). Microplastics, particles smaller than 5 mm, have been detected in various aquatic environments and organisms, including the wharf roach (*Ligia* spp., Subphylum Crustacea and Order Isopoda) (Bajt, 2021; Choi et al., 2023).

Polystyrene is a versatile plastic widely used in packaging and other industries. Expanded polystyrene (EPS) is commonly employed in the fishing industry. Wind and ocean currents transport and accumulate debris in coastal areas (Kuroda et al., 2024; Sagawa et al., 2018). A field survey demonstrated the prevalence of EPS in wharf roaches collected from rocky shorelines in Japan (Lee et al., 2025).

Microplastics can cause infertility, oxidative stress, inflammation, immunotoxicity, neurotoxicity, and metabolic disruption in aquatic organisms, including fish, crabs, annelids, and mollusks (Hodkovicova et al., 2022). Moreover, they can disrupt the gut microbiota, leading to dysbiosis and microbial community imbalance, adversely affecting health (Fackelmann and Sommer, 2019; Shi et al., 2024). Tamura et al. (2024) observed altered shoaling behavior in medaka fish exposed to 2 µm polystyrene microplastics and attributed this finding to the dysbiosis of bacteria producing short-chain fatty acids and subsequent changes in neurotransmitter levels. Intestinal gene expression is also altered in fish exposed to microplastics (Assas et al., 2020; Zhang et al., 2023). Although microplastics have been suspected to affect gut health, their impact on organisms inhabiting rocky seashores remains unclear.

The size of polystyrene microplastics significantly influences their impact on the gut microbiome, with small particles exerting more substantial effects on the bacterial and fungal composition and metabolic pathways than large particles (Gao et al., 2023). Isopods have diverse gut microbiomes (Bouchon et al., 2016). Exposure to polystyrene microplastics has been associated with changes in the core microbial population structure and expression of genes associated with antioxidative, detoxification, and immune system functions (Wang et al., 2021). Similar effects may be observed in other species. Heffner et al., (2023) indicated that soil-dwelling isopods enhance the interactions between bacteria and fungi involved in methane processes. Furthermore, Lee et al., (2024) examined the gut microbiome composition of *Ligia* spp. collected from areas contaminated with EPS.

Wharf roaches frequently ingest expanded polystyrene (EPS) in sandy and rocky coastal environments and fragment it (Lee et al., 2025). This frequent and species-specific ingestion suggests that wharf roaches may not only consume EPS incidentally but also utilize it as a food source, thereby possibly contributing to the degradation of EPS microplastics through fecal pellets. However, the effects of EPS ingestion on gene expression and gut microbial communities in wharf roaches remain unclear. Thus, this study aimed to investigate the effects of EPS on gene expression and gut microbiome composition in wharf roaches. The results of this study may serve as a basis for (Hoyer and Hyttinen, 2009) elucidating the impact of EPS on organisms inhabiting rocky shoreline habitats that may be exposed to and consume plastics.

## 2 Materials and methods

### 2.1 Materials

An EPS board was purchased from a hardware store (NAFCO, Motooka, Nishi Ward, Fukuoka, Japan). Aquarium gravel obtained from Pets Value Co., Ltd. (Hyogo, Japan) was used in wharf roach rearing tanks. Artificial seawater was prepared using Marine ART SF-1 (Tomita, Tokushima, Japan).

### 2.2 Sample collection and maintenance

Metagenomic and mRNA-seq analyses were performed to investigate the effects of EPS ingestion on gene expression and gut microbiota in wharf roaches. Wharf roaches were collected from Nishinoura fishing ports in Nishi-Ku, Fukuoka Prefecture, Japan, in April 2021. Approximately 30 collected wharf roaches were placed in a glass container (25 cm high, 35 cm wide, and 21.5 cm long) prepared with artificial seawater in a Petri dish (diameter: 17 cm) and a layer of pebbles (height: 1 cm). They were acclimatized to laboratory conditions for a week, during which dead individuals were promptly removed to prevent cannibalism. The seawater was changed every 2 days, and frozen brine shrimp (Vitacrine Baby Brine, Hikari; Kyorin Co., Ltd., Hyogo, Japan) was provided every 3 days.

### 2.3 Expanded polystyrene feeding test for mRNA-seq and microbiome analyses

After the acclimation period, 10 wharf roaches were randomly selected and individually transferred into cubic plastic containers (12 cm high, 21 cm wide, and 13.5 cm long), which were also set up with a layer of pebbles in a Petri dish (diameter: 5 mm) filled with artificial seawater. The experimental wharf roaches were starved for 3 days to induce a fasting state before exposure to EPS. Following this, they were divided into two groups: a control group that continued fasting and an EPS feeding group that was provided only EPS blocks (approximately 50 mg) as food. The experiment was conducted for a week, after which the wharf roaches were dissected to extract their guts, and total RNA and DNA were isolated using TRIzol Reagent (Thermo Fisher Scientific, Tokyo, Japan). Individuals carrying eggs were excluded from this study. The extracted DNA and RNA were sequenced using a NovaSeq 6000 system (Illumina, Inc., San Diego, CA, USA).

### 2.4 RNA-seq analysis

Raw reads of the mRNA-seq data were processed using fastq for quality control purposes. The analysis was conducted using a de novo approach in Trinity, with only clean data (contig BUSCO score, 97.7%). These contigs were used for downstream analyses, including gene expression quantification and normalization using edgeR (https://bioconductor.org/packages/release/bioc/html/edgeR.html). Differentially expressed genes (DEGs) were identified based on statistical thresholds (p < 0.01 and false discovery rate (FDR) < 0.01). Heatmaps were generated to visualize the expression patterns of the DEGs across the samples. Multidimensional scaling (MDS) was also performed. The contigs were annotated using BLAST searches against public databases to identify the putative gene functions. Functional categories associated with the DEGs (∣logFC∣>2) were explored to facilitate the interpretation of the biological implications of differential expression. Genes were annotated using the *Arthropoda* database (https://www.ncbi.nlm.nih.gov/datasets/taxonomy/6656/).

### 2.5 Metagenome analysis on wharf roaches fed with EPS

#### 2.5.1 Metagenome analysis

Whole-genome analysis of the gut feces from the wharf roaches was conducted using a NovaSeq 6000 system. Based on the acquired raw data, classification into the categories of bacteria, archaea, eukaryotes, and viruses, and their relative proportions was conducted. Based on the relative proportion data, analysis by non-metric multidimensional scaling (NMDS) was performed using the “vegan” library of the R software and visualized using the “ggplot2” library. Statistical evaluation was performed using the pairwise Adonis library. All genomic datasets were deposited in the GenBank Sequence Read Archive database.

#### 2.5.2 Correlation analysis

The biomes in the feces of *Ligia* spp. were classified through correlation analysis. A correlation network was visualized after calculating the Pearson correlation coefficient (|r|>0.7) using R software, as previously described. The modularity class was automatically discerned using Gephi (version 0.10.1) (https://gephi.org) and visualized (Force Atlas with Noverlap).

#### 2.5.3 Feature selection

Random forest (Breiman, 2001), a supervised machine learning (ML) algorithm with bagging (bootstrap aggregating) (Miyamoto and Kikuchi, 2023), was used to extract the feature components in the dataset. The feature components were selected using the “randomForest” package in R software (https://cran.r-project.org/web/packages/randomForest/index.html).

#### 2.5.4 Inference of common factors across different farms

In brief, important factors were searched using exploratory factor analysis (EFA) (Peprah et al., 2021; Schmitt, 2011) in the “psych” and “GPArotation” library packages in R software. The analysis codes were obtained from the website (https://cran.r-project.org/web/packages/psych/index.html). The number of factors was calculated using the “VSS” function for the very simple structure and the “fa” function as previously described. EFA was calculated using the minimum residual (minres) method. The function “promax,” an oblique rotation that allows factors to be correlated, was used as the calculation method instead of an orthogonal rotation. The other values were as follows: h2, communality score; u2, uniqueness score; and com, complexity, an information score that is generally related to uniqueness. These selected factors calculated by the “fa” function were visualized using the “heatmaply” library package in R software (https://cran.r-project.org/web/packages/heatmaply/vignettes/heatmaply.html).

#### 2.5.5 Structural equation modeling

Structural equation modeling (SEM) for confirmatory factor analysis (CFA) was performed using the “lavaan” package in R software (Rosseel, 2012; Rosseel and Loh, 2022) as previously described (Miyamoto et al., 2023, 2022; Tamura et al., 2024). The analysis code was based on the website (https://lavaan.ugent.be). As CFA requires a hypothesis, the bacteria selected in the association, linear discriminant, and energy landscape analyses were used as factors for latent components of the data. The hypothesized model was statistically estimated using maximum likelihood parameter estimation with bootstrapping (n = 1000) using the “lavaan” and “semsem” functions. The goodness-of-fit of the model was evaluated using the chi-square p-value (p > 0.05, not significant), degrees of freedom (df), comparative fit index (cfi) (>0.95), Tucker–Lewis index (tli) (>0.95), normed fit index (nfi) (>0.95), relative fit index (rfi) (>0.95), root mean square error of approximation (rmsea) (<0.05), and standardized mean residual (srmr) (<0.08), goodness-of-fit index (gfi), and adjusted goodness-of-fit index (agfi) (Hooper et al., 2008). Candidate models with lower Akaike information criterion values and smaller differences between the gfi and agfi (>0.95) were selected. Path diagrams for the superior models were visualized using the “semPlot” package in R (Epskamp et al., 2019).

#### 2.5.6 Statistical causal inference

Three types of causal inferences were conducted with some modifications, as reported previously. Briefly, causal mediation analysis was performed using the “mediation” (Tingley et al., 2014) package in R software based on a tutorial website (https://rpubs.com/Momen/485122). First, each regression relationship with “∼” in the selected model was assessed using the “lm” function. Subsequently, the values of the causal relationships between bacterial candidates as mediators and outcomes were evaluated using the “mediation” function. The estimated average causal mediation effect (ACME), average direct effect (ADE), and proportion of the total effect through mediation were calculated by nonparametric bootstrapping (“boot=TRUE”) with “sims=1000” as the number of iterative calculations and a quasi-Bayesian approximation (“boot=FALSE” as a command). The computable data are listed in Table S6. A Bayesian score-based approach (BayesLiNGAM) (Hoyer and Hyttinen, 2009) that takes advantage of non-Gaussian distributions when evaluating linear acyclic causal models was applied to the selected groups related to SEM-selected components. BayesLiNGAM was established using the “fastICA” package in R software based on the information from the specialized website (https://www.cs.helsinki.fi/group/neuroinf/lingam/bayeslingam/). The “fastICA” package in R was used according to the other information on the website (https://cran.r-project.org/web/packages/fastICA). The seasonal directed acyclic graphs calculated by BayesLiNGAM were visualized as networks in the “igraph” package in R.

### 2.6 Other statistics

The F-test and Shapiro–Wilk tests were used to determine equal variances, Gaussian distributions, or non-Gaussian distributions. The Shapiro–Wilk test was also used in the “shapiro.test” function of R software. The unpaired t-test and Mann–Whitney U test were performed as appropriate, depending on the data type. Statistical significance was set at p < 0.05, and a tendency was assumed at p < 0.1. These calculation data were prepared using Microsoft 365 and R software (versions 4.3.0 and 4.3.3). Data are presented as mean ± SE.

## 3. Results

### 3.1 Expanded polystyrene feeding test for mRNA-seq analysis

The transcripts obtained from each treatment were analyzed using MDS plots. The filtered sample count data were normalized for size between libraries, and an MDS plot was used to assess the integrity of the sample transcript counts. The MDS plot (Fig. 1) shows the differences between the EPS-fed and control groups. Samples from the EPS-fed group clustered in the upper right quadrant, whereas those from the control group were positioned in the lower left quadrant. The first two principal components accounted for 25% and 20% of the total variance, respectively.

**Fig. 1.**
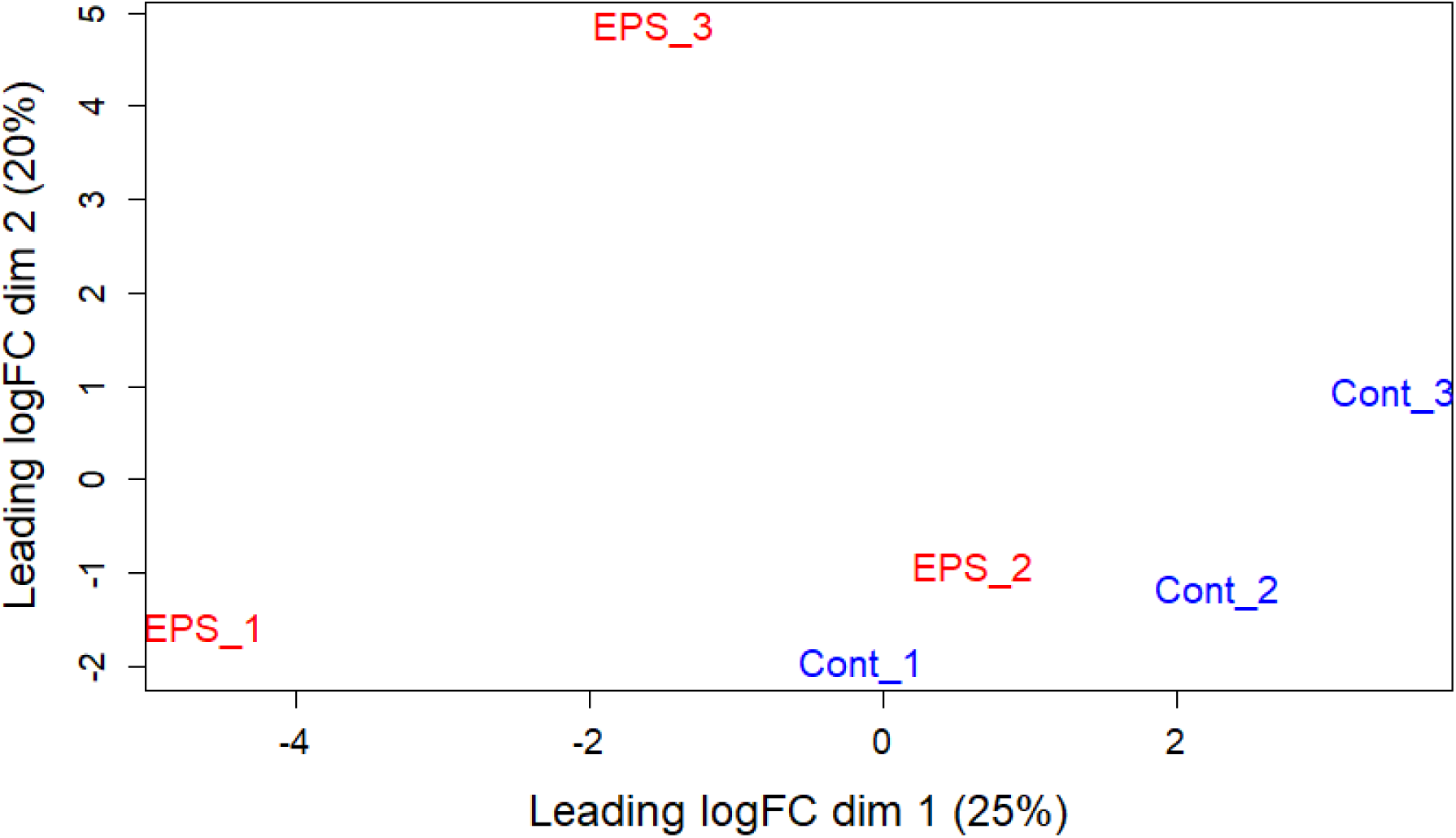
A multidimensional scaling (MDS) plot illustrating the similarity in response among wharf roaches (*Ligia* spp.) specimens based on similarities in the observed transcript counts between the qualified sequence read libraries. EPS_1, EPS_2, and EPS_3: expanded polystyrene (EPS)-exposed specimens; Con_1, Con_2, and Con_3: control

RNA-seq data analysis revealed substantial alterations in gene expression following exposure to EPS. Utilizing a threshold of |log FC| ≥ 2, we identified 182 genes with significantly increased expression and 267 genes with significantly decreased expression in the EPS-fed group compared with the control group (Table 1, Table S1). Fig. 2 presents a heat map of the 267 DEGs that satisfied the criteria of p < 0.01, FDR < 0.05, and ∣logFC∣>2. Visualization revealed distinct differences in the clustering patterns between the EPS-exposed and control groups. Heat map analysis indicated that the gut of the wharf roaches in the EPS-fed group exhibited significant alterations in global gene expression compared with that of the wharf roaches in the control group.

**Fig. 2.**
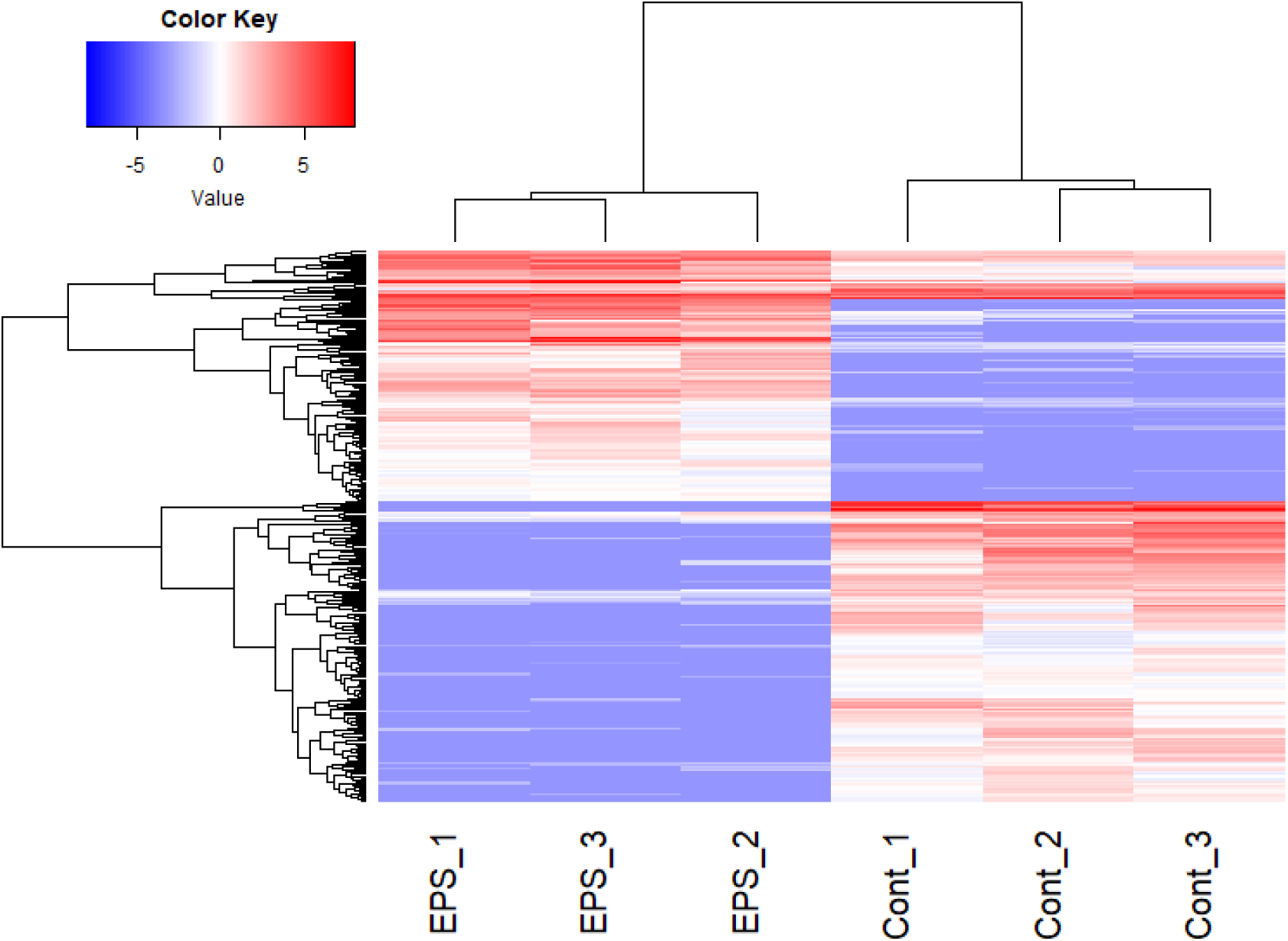
Heatmap of differentially expressed genes (DEGs) in the gut of wharf roach fed EPS. Abbreviations: EPS_1, EPS_2, and EPS_3, EPS-fed groups; and Con_1, Con_2, and Con_3, control groups without EPS treatment.

**Table 1.** List of remarkable genes related to xenobiotic metabolism in wharf roach (*Ligia* spp.) that significantly changed in expression after EPS feeding (p < 0.01, FDR < 0.05, and ∣logFC∣>2).

| Seq-name | Description | logFC | p-value | FDR |
| --- | --- | --- | --- | --- |
| TRINITY_DN9248_c0_g1_i12 | Glycosyltransferase-like domain-containing protein 1-like [*Armadillidium nasatum*] | 7.462 | 1.72E-05 | .005 |
| TRINITY_DN105608_c0_g1_i1 | Cytochrome oxidase subunit I, partial [*Aegla franciscana*] | 5.323 | 2.63E-06 | .001 |
| TRINITY_DN4368_c0_g1_i12 | Betaine--homocysteine S-methyltransferase 1-like [*Penaeus monodon*] | 4.749 | 1.55E-05 | .004 |
| TRINITY_DN68926_c0_g1_i13 | Cytochrome P450 302a1, mitochondrial-like [*Penaeus monodon*] | 4.574 | 4.77E-05 | .010 |
| TRINITY_DN110_c0_g1_i3 | 2-Acylglycerol O-acyltransferase 1-like [*Eriocheir sinensis*] | 4.381 | 0.0003 | .045 |
| TRINITY_DN420_c1_g1_i1 | Cytochrome P450 [*Armadillidium nasatum*] | 4.315 | 8.61E-05 | .016 |
| TRINITY_DN1408_c0_g1_i1 | Pyridoxine-5'-phosphate oxidase [Armadillidium vulgare] | 3.817 | 0.0003 | .042 |
| TRINITY_DN81510_c2_g1_i1 | Betaine--homocysteine S-methyltransferase 1-like [Penaeus monodon] | -3.832 | 0.0003 | .040 |
| TRINITY_DN12448_c0_g1_i1 | Lysophospholipid acyltransferase 7 [Armadillidium nasatum] | -4.303 | 0.0001 | .028 |
| TRINITY_DN16_c1_g1_i3 | Putative ZDHHC-type palmitoyltransferase 6 [Armadillidium nasatum] | -4.383 | 8.96E-05 | .016 |
| TRINITY_DN1747_c0_g1_i13 | Galactose-1-phosphate uridylyltransferase [Armadillidium nasatum] | -4.451 | 6.36E-05 | .013 |
| TRINITY_DN3342_c2_g1_i6 | Cytochrome P450 9e2, partial [Armadillidium vulgare] | -5.237 | 2.85E-05 | .007 |
| TRINITY_DN11586_c0_g1_i35 | Histone acetyltransferase KAT7, partial [Armadillidium nasatum] | -6.159 | 1.81E-05 | .005 |
| TRINITY_DN21794_c0_g1_i6 | N-acetyltransferase ESCO2 [Armadillidium nasatum] | -6.508 | 0.0003 | .040 |
| TRINITY_DN8169_c0_g1_i14 | Protein-lysine N-methyltransferase n6amt2 [Armadillidium nasatum] | -6.778 | 0.0001 | .021 |
| TRINITY_DN10879_c0_g1_i2 | 5-Methylcytosine rRNA methyltransferase NSUN4-like isoform X1 [Eriocheir sinensis] | -7.252 | 3.33E-05 | .008 |
| TRINITY_DN2539_c0_g1_i1 | Peroxidase skpo-1 [Armadillidium vulgare] | -7.584 | 1.16E-05 | .003 |
| TRINITY_DN768_c1_g1_i1 | KAB7495655.1 tRNA methyltransferase [Armadillidium nasatum] | -8.096 | 2.22E-06 | .0009 |
| TRINITY_DN1101_c1_g1_i19 | Protein-cysteine N-palmitoyltransferase HHAT-like [Penaeus japonicus] | -8.995 | 1.20E-07 | .83E-05 |
| TRINITY_DN1416_c0_g1_i19 | Glutathione peroxidase-like isoform X1 [Cherax quadricarinatus] | -10.429 | 9.96E-10 | .67E-06 |

Approximately 81% of these genes were annotated using the Arthropoda database. Transcriptional analysis revealed significant EPS-induced changes in the expression of genes related to detoxification, metabolism, and cell stability. Genes associated with xenobiotic metabolism, such as members of the cytochrome P450 family (cytochrome P450 302a1; logFC, 4.6, cytochrome P450; logFC, 4.3, and cytochrome P450 9e2; logFC, −5.2), showed significantly altered expression levels, highlighting their potential roles in processing and detoxifying harmful substances. Similarly, genes involved in glycosylation and triglyceride synthesis, including the glycosyltransferase-like domain-containing gene (logFC = 7.5) and 2-acylglycerol O-acyltransferase gene (logFC = 4.4), were expressed differently. The expression levels of genes regulating cellular processes, such as histone acetyltransferase KAT7 (logFC, −6.2) and N-acetyltransferase ESCO2 (logFC, −6.5), were downregulated, suggesting potential effects on gene regulation and chromosomal cohesion. Additionally, genes related to methylation balance (betaine-homocysteine S-methyltransferase; logFC, 4.7), vitamin B6 activation (pyridoxine-5’-phosphate oxidase; logFC, 3.8), antioxidant defense (glutathione peroxidase-like isoform; logFC, −10.4), and protein translation accuracy (tRNA methyltransferase; logFC, −8.1) were differentially expressed, indicating their involvement in the response to EPS exposure. In particular, the involvement of the p53 signaling pathway was highlighted. NUP53 exhibited a log fold change (logFC) of −5.52, indicating strong downregulation.

### 3.2. Expanded polystyrene feeding test for microbiome analysis

Based on genomic data, categorical diversity was evaluated using NMDS. No statistically significant differences were observed (Fig. S1). Therefore, as a relative quantitative assessment of biota, we compared groups of organisms with abundance ratios ≥1%. *Escherichia* was detected in the control and EPS treatment groups at an abundance ratio of ≥10% (Fig. 3a). The abundance of *Pseudomonas* was lower in the EPS group than in the control group (p = 0.078). As for the archaeal flora, *Halorhabdus* and *Natrialba* had ratios ≥10% between the control and EPS groups (Fig. 3b). The abundance of *Halorubrum* was lower in the EPS group than in the control group (p = 0.1). The abundances of *Halomicrobium* and *Haloquadratum* were higher in the EPS group than in the control group (p = 0.071; p = 0.022). Among the eukaryotes, *Dictyostelium*, *Loa*, and *Hydra* were detected at ratios ≥10% in the control and EPS groups (Fig. 4a). The abundances of *Loa* and *Volvox* were higher in the EPS group than in the control group (p = 0.091; p = 0.034). Meanwhile, the abundance of *Gallus* was lower in the EPS group than in the control group (p = 0.084). Furthermore, *Chlorovirus, Rhadinovirus, Ichinovirus,* and *Macavirus* had abundance ratios ≥10% between the control and EPS groups (Fig. 4b). The abundances of *Ichinovirus* and *Varicellovirus* were higher in the EPS group than in the control group (p = 0.1 and p = 0.001, respectively).

**Fig. 3.**
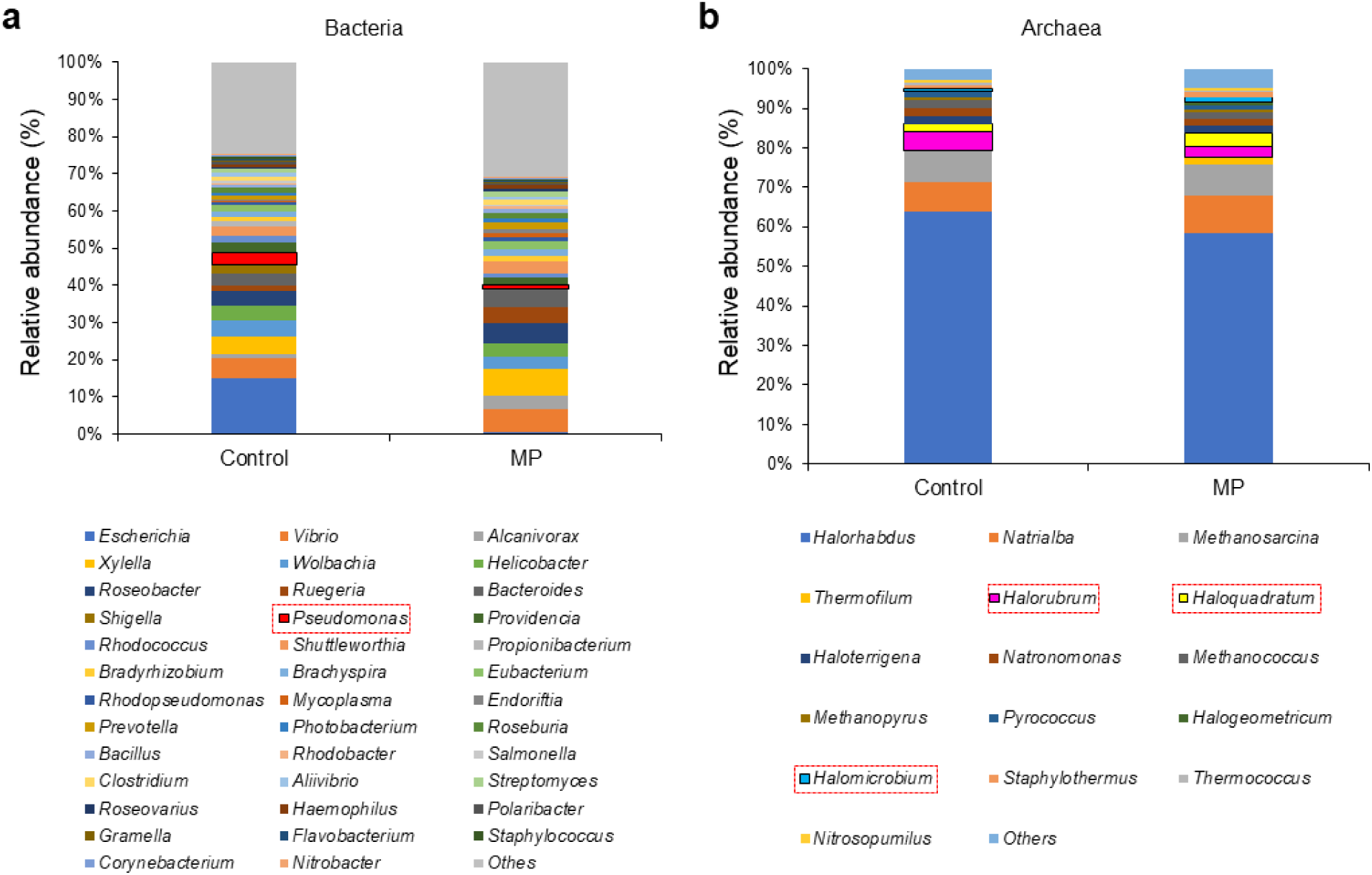
Relative abundance of bacterial and archaeal populations as symbiotes in the wharf roach. The majority of the populations of (a) bacteria and (b) archaea were visualized (>1% of the total population in each category). The dotted squares show the components of the statistical features (p < 0.1). Abbreviations: Cont, control group without EPS treatment; EPS, group with EPS treatment.

**Fig. 4.**
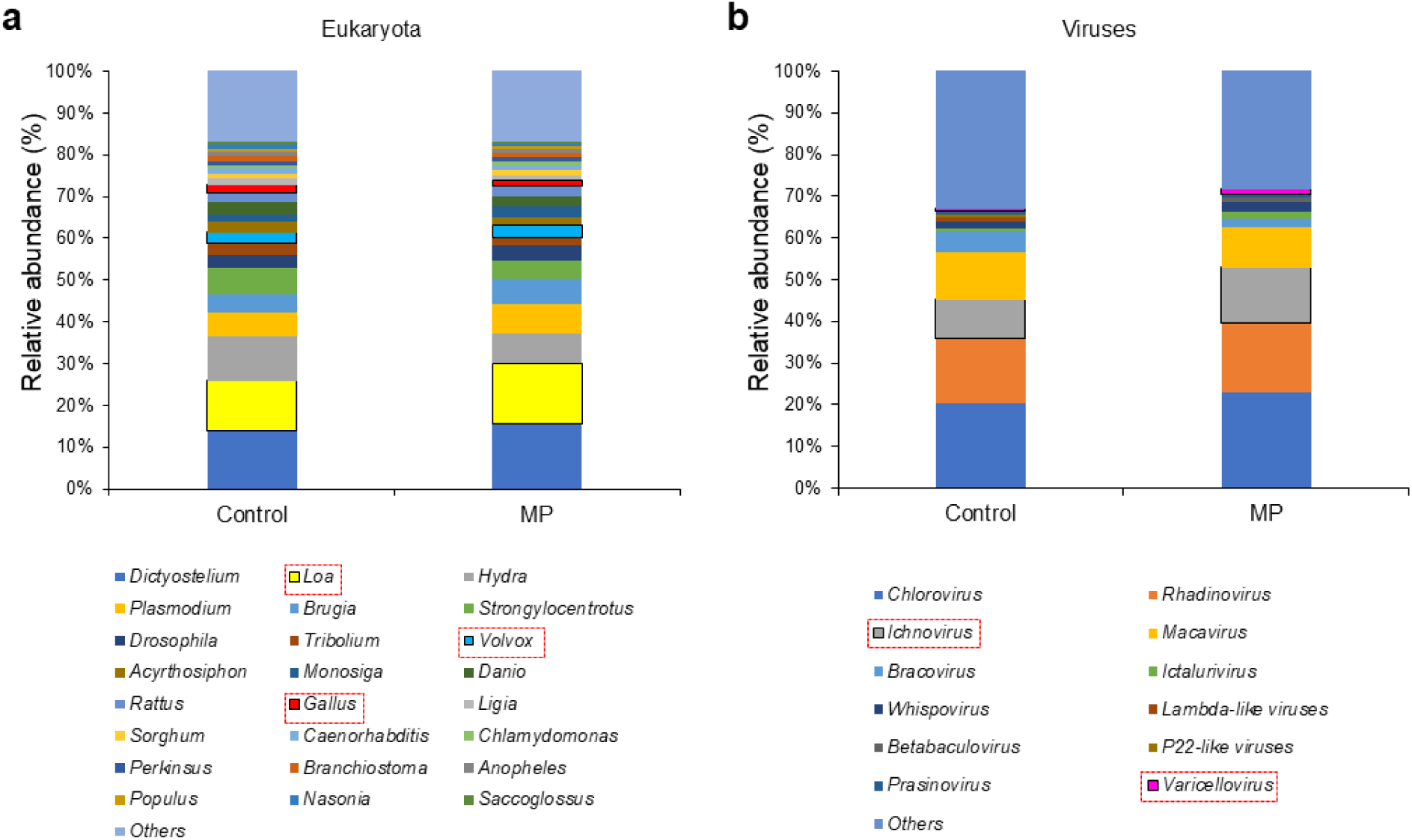
Relative abundance of eukaryotic and viral populations as symbiotes in the wharf roach The majority of the populations of (a) eukaryotes and (b) viruses were visualized (>1% of the total population of each category). The dotted squares indicate components with statistical features (p < 0.1). Abbreviations: Cont, control group without EPS treatment; EPS, group with EPS treatment.

### 3.3 Correlation network of predominant factors

Categorical correlation analyses were performed for factors that showed marked differences (p < 0.2) between the control and EPS treatment groups. Correlation networks (>0.1% relative abundance and |r|>0.7) are shown. In the positive correlation group, bacteria *Geobacter* and *Spirosoma*; archaea *Dusulfurococcus*, *Haloquadratum*, and *Halomicrobium*; eukaryotes *Halalkalicoccus*, *Methanospirillum*, *Arabidopsis*, *Loa*, and *Volvox*; and viruses *Ichnovirus*, *T4-like viruses,* and *Varicellovirus* formed a network (Fig. 5a). In the negative correlation group, bacteria *Congregibacter* and *Pseudomonas* and eukaryotes *Apis*, *Caenorhabditis*, *Magnaorthe*, and *Trichoplax* formed a network (Fig. 5b).

**Fig. 5.**
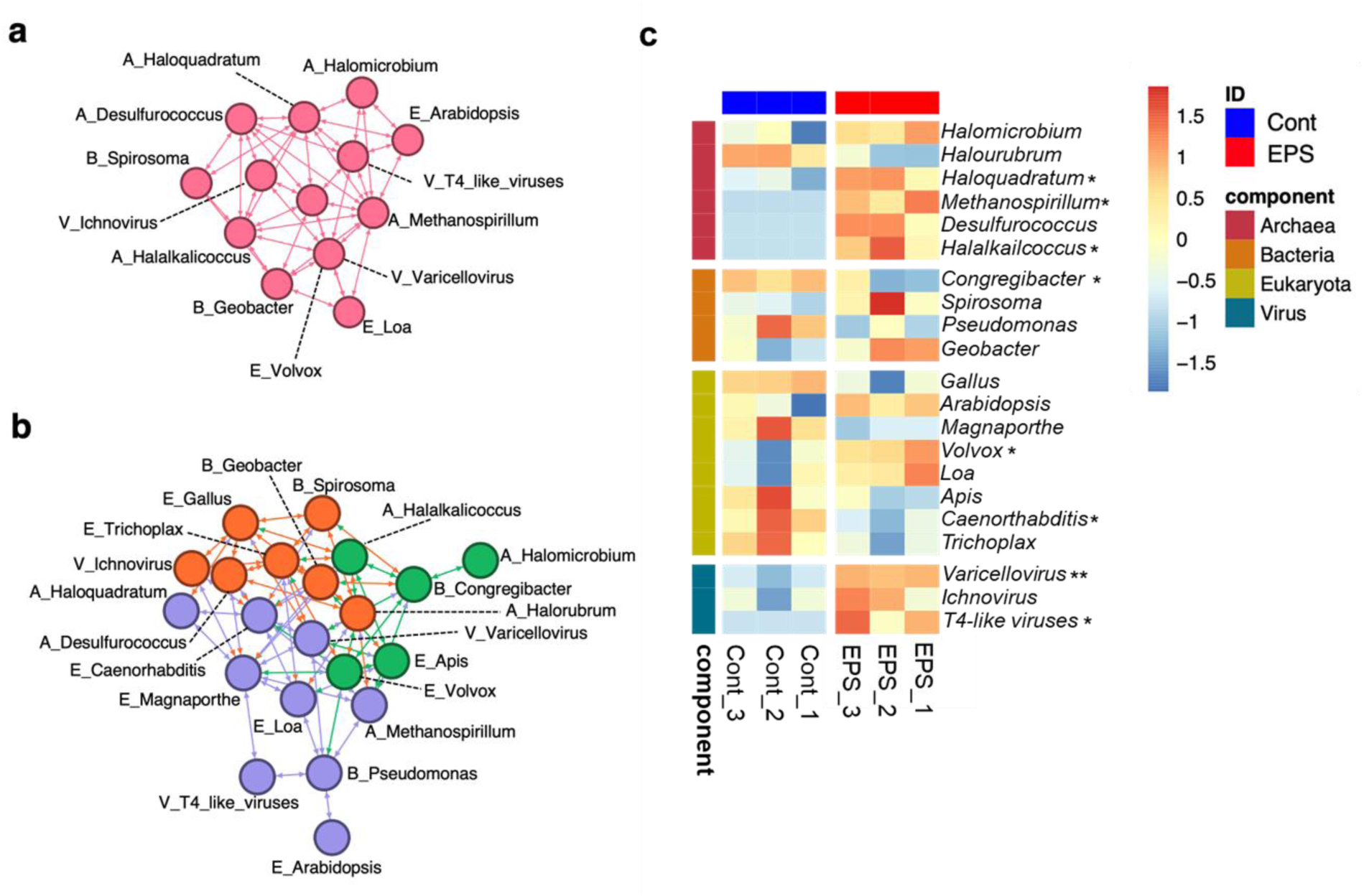
Correlation analysis of the analysis-classified symbiotes in the wharf roach. A correlation network of the symbiotes in the wharf roach is shown. The criteria for the correlation analysis were set as |r| > 0.7. (a) A positive and (b) a negative correlation cluster was visualized with the color-classified modularity class calculated by Gephi 0.10 (https://gephi.org). (c) Heatmap showing the relative abundance of these components depending on the conditions. Abbreviations: A_, archaea; B_, bacteria; E_, eukaryote; V_, virus; Cont, group without EPS treatment; and EPS, group with EPS treatments. Asterisks indicate significance (* p < 0.05; ** p < 0.01).

The abundance ratios of these factors were compared between the two groups (Fig. 5c and Table S2). In particular, those of bacteria *Congregibacter*; archaea *Haloquadratum*, *Halalkalicoccus*, and *Methanospirillum*; eukaryotes *Caenorhabditis* and *Volvox*; and viruses *T4-like viruses* and *Varicellovirus* were significantly dominant (p < 0.05).

These results suggest that the symbiotes of *Ligia* spp. are linked to factors beyond the biota. Considering these results, we proceeded to the factor analysis and prediction of the causal structure.

### 3.4 Classification of important factors based on ML and EFA

As a result of generating features using random forest, which is a type of ML, for the factor groups classified by correlation analysis, the numerical values of each did not differ by more than twofold (Fig. S2), and these differences were not always statistically significant (Table S2). However, these factors showed marked differences (p ≤ 0.1) between the control and EPS groups in this study. Therefore, we investigated the relationships between the feature factors using factor analysis. First, the optimal number of factors was determined using a scree plot (Fig. S3a). The value of Cattell scree was two as the factor number, and the value of Kaiser-Guttman was six, but the value of VSS (very simple structure) was four (Revelle and Rocklin, 1979; Taguchi et al., 2024). The tendency to adopt VSS-calculated values has recently been reported; therefore, the VSS value was used for subsequent calculations. When a correlation was present between these factors (Figs. 5a and 5b), the varimax rotation could not be applied in the formula “rotate”; thus, promax rotation was adopted, and factor analysis was performed (Fig. S3b). The value of h2 indicates that the cooperation (Table S3) was not low (h2 > 0.3). Calculation results showed that these factor groups worked in concert rather than as individual factors. Therefore, to predict the most important factor groups, we constructed a phylogenetic tree of these groups using heatmaps. *T4-like viruses*, *Varicellovirus, vanadate*, and *Methanospirillum* were depicted as more closely clustered in proximity to EPS administration (Fig. 6a). Therefore, these factors were calculated using a covariance structural analysis. Consequently, we generated a group of structural equation models with optimal values (Fig. 6b, Table S4). *T4-like viruses* and *Methanospirillum* were positively correlated, with a correlation coefficient of 0.93 (p < 0.001). *Methanospirillum* and EPS administration were positively correlated, with a contribution value of 0.96 (p < 0.001). *Methanospirillum* and *Varicellovirus* were positively correlated, with a contribution value of 0.08 (p = 0.791). *Varicellovirus* and EPS administration (Tst) were positively correlated, with a contribution value of 0.91 (p = 0.001). Mediated analyses of these factors (Table S5) revealed the significance of the factor groups. However, the direct effect of a single factor (ADE) and the mediated effect of two specific factors (ACME) could not be calculated. Therefore, these groups of factors were deemed important for the overall structure of a group (Fig. 6b).

**Fig. 6.**
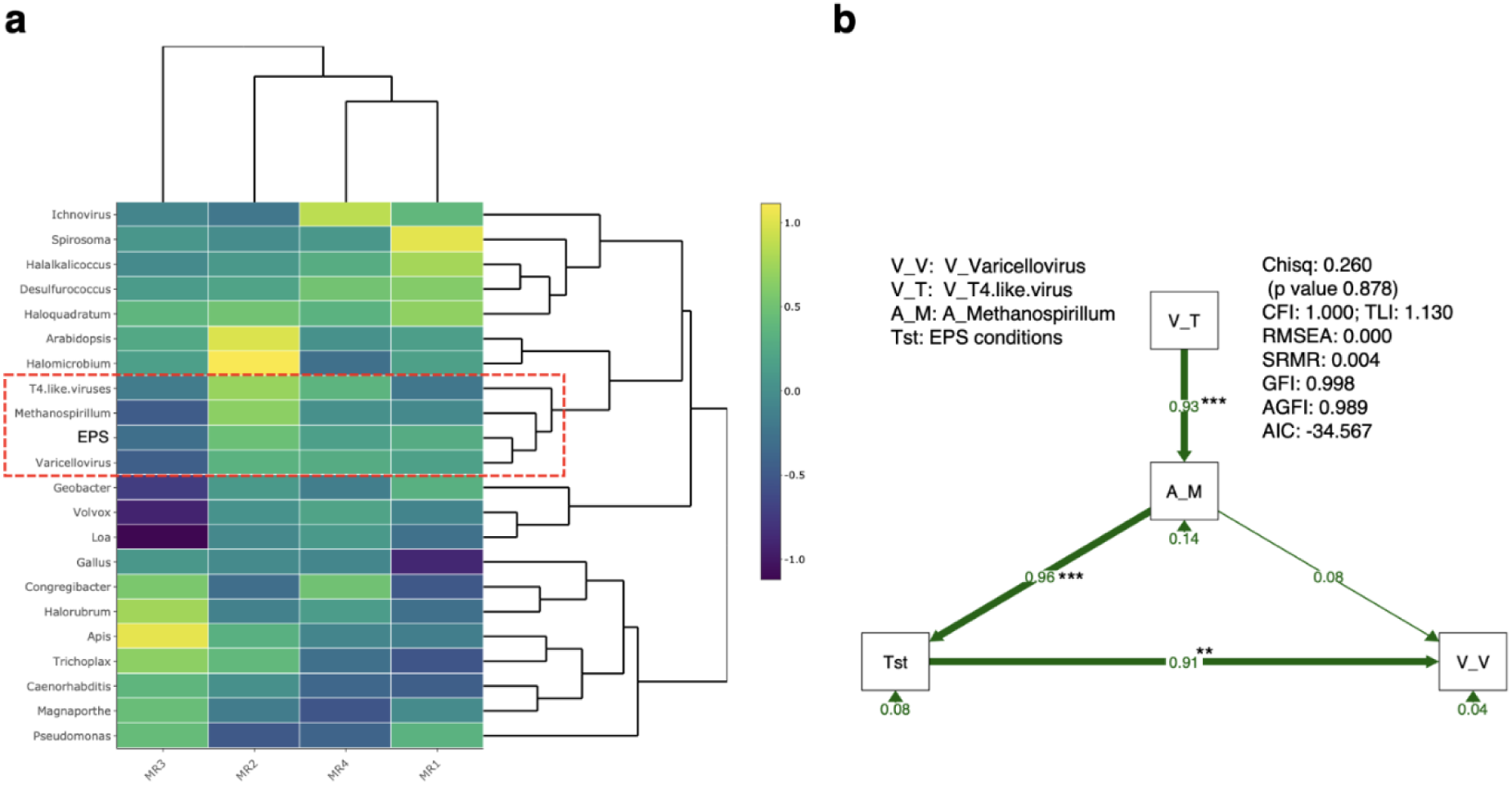
Classification of features of the symbiote composition by EFA. (a) Heatmap of components obtained from EFA setting 4 as the factor number. MR indicates the minimal residual method. The right bar indicates the extent of the contribution. Components with commonality scores (>0.3 as h2) are displayed. h2 shows the commonality score, which is the amount of variance in the variables explained by the factors. The components within the red square are strongly associated with EPS treatment (EPS). (b) Optimal structural equation model based on the components selected in Fig. 6a. Abbreviations: Tst, EPS treatment; A_M, A_Methanospirillum; V_T, V_T4.like.virus; V_V, V_Varicellovirus. The main fit indices of the models are shown in the figure, and the other indices are listed in Table S4 (Supporting Information). Abbreviations: chisq, Chi-square: χ2; p value, p values (Chi-square); CFI, comparative fix index; TLI, Tucker–Lewis index; RMSEA, root mean square error of approximation; SRMR, standardized root mean residuals; GFI, goodness-of-fit index; and AGFI, adjusted goodness-of-fit index. *, p < 0.05; and ***, p < 0.001.

## 4. Discussion

Our results clearly demonstrate that EPS consumption by wharf roaches in rocky coastal environments induces gene responses and disturbs microbial equilibrium in the digestive system. A feeding experiment validated EPS ingestion and identified alterations in the expression of genes associated with xenobiotic metabolism, glycosylation, cell processing, and dysbiosis in the intestinal tract of the wharf roaches. The populations of *Methanospirillum* spp., which produce methane, and viruses significantly increased in the EPS group. These results indicate that EPS consumption can alter gene expression, disrupt microbial balance, and potentially contribute to global warming and climate change.

The upregulated expression of cytochrome P450 genes underscores their critical role in detoxifying harmful substances during EPS exposure. These enzymes possibly mitigate oxidative damage by metabolizing toxic compounds, consistent with their established functions in xenobiotic metabolism under environmental stress (Regoli and Giuliani, 2014). Similarly, the upregulated expression of glycosyltransferase-like domain-containing proteins highlights the importance of glycosylation in maintaining membrane integrity and enabling adaptive stress responses. Glycosylation supports cellular communication and signaling, which are essential for mitigating environmental stress and facilitating recovery (Zhu et al., 2019). The upregulated expression of betaine-homocysteine S-methyltransferase (BHMT1) suggests its role in regulating methylation balance and metabolic flexibility. BHMT1, particularly in crustaceans, is associated with osmoregulation and stress-related hormone synthesis, which are critical for genomic stability and adaptive responses (Yin et al., 2025).

By contrast, the downregulated expression of the glutathione peroxidase-like isoform indicates a potential weakening of the antioxidant defenses following EPS exposure. This enzyme plays a key role in mitigating oxidative stress by neutralizing the peroxides. Reduced expression may increase cellular vulnerability to reactive oxygen species, thereby increasing the risk of oxidative damage and cellular dysfunction.

Genes involved in chromatin remodeling, such as ESCO2 and KAT7, exhibited significant transcriptional changes. ESCO2 facilitates cohesin-mediated chromatin stability during DNA replication, whereas KAT7 supports transcription by loosening chromatin through histone acetylation. Alterations in their expression suggest that EPS exposure can destabilize chromatin organization, impairing transcriptional regulation and epigenetic stability. These disruptions can lead to long-term genomic instability and an increased risk of maladaptive responses. Previous studies have also supported these findings. For instance, ESCO2 depletion destabilizes cohesin and impairs transcriptional regulation (Fu et al., 2023; Nishiyama et al., 2010), whereas KAT7 dysfunction is linked to epigenetic misregulation under environmental stress (Yang et al., 2022). Similarly, xenobiotic exposure in zebrafish alters the expression of detoxification-related genes (Chen et al., 2017), and microplastics contaminated with persistent organic pollutants affect the expression of vitellogenin and choriogenin in medaka (Rochman et al., 2014).

The present study aligns with previous transcriptome analyses of aquatic organisms, showing that plastic significantly influences immune responses, oxidative stress, and metabolic pathways (Kazmi et al., 2024). Additionally, the differential expression of immune and oxidative stress-related genes, such as hemocyanin, supports the activation of stress-related signaling pathways in *Ligia exotica* exposed to EPS. Notably, the present study used a controlled laboratory setting (EPS-only diet and fasting), which resulted in more pronounced heterologous expression of the target gene changes in gene expression than in field studies (Choi et al., 2023). These findings emphasize the importance of understanding transcriptomic changes under various environmental conditions to elucidate the long-term effects of EPS exposure.

The increase in virus-related transcripts, such as autophagy-related protein 101-like and influenza virus NS1A-binding protein homolog isoform X2 (Table S5.1) (Zhirnov and Klenk, 2013), supports the strong positive correlation observed between the ingestion of EPS and the abundance of viruses such as T4-like virus and *Varicellovirus* in the gut microbiota analysis (Fig. 5b).

The observed increase in *Methanospirillum* abundance in the EPS-fed group was noteworthy. *Methanospirillum* belongs to Class 2 methanogenic archaea, which utilize a broader range of carbon sources than Class 1. This finding suggests that the consumed EPS may have served as a carbon source for *Methanospirillum*. Furthermore, methanogenic archaea and viruses reportedly interact and exhibit ecological associations (Medvedeva et al., 2023; Ngo et al., 2022), supporting the observed positive correlation between the increase in *Methanospirillum* and the abundance of viruses in the gut of wharf roaches (Fig. 6b). Other studies have reported that the presence of plastics under anaerobic digestion conditions in wastewater treatment facilities can enhance methane production (Quecholac-Piña et al., 2020). Based on these findings, we hypothesized that EPS in the digestive tract of *L. exotica* contributes to the creation of a more anaerobic gut environment, activating methanogenic archaea during its degradation. Moreover, previous research indicated increased hydrogen compound production when plastics are included in the diet of terrestrial isopods (*Porcellio scaber*) (Hink et al., 2023), which are ecologically closely related to wharf roaches. This finding supports the observed increase in *Methanospirillum* populations in the present study. Mahane is a major contributor to global warming. The potential increase in methanogenic archaea due to plastic ingestion by organisms suggests that these processes contribute to methane production. Previous studies have reported that plastic ingestion by zooplankton accelerates ocean deoxygenation. Thus, plastic pollution affects not only organisms but also the global environment, highlighting its far-reaching consequences. However, considering that EPS contains various additives, further research is necessary to determine whether the methanogenic archaea in the gut of the wharf roach utilizes the plastic polymer or the additives as carbon sources. Further studies are required to quantify the increase in methane production resulting from plastic ingestion.

In the present study, gut microbiota analysis of wharf roaches exposed to EPS for 10 days revealed dysbiosis. The effects of EPS administration on the gut microbiota were investigated using ML and structural equations. The rationale for establishing the control group as a starving group in this experiment was twofold: to ascertain the digestibility of EPS by wharf roaches and to minimize alterations in the gut microbiota of wild-caught wharf roaches used for arbitrary food provisioning.

Wharf roaches have been utilized as a model organism for monitoring coastal pollution in several studies, but mostly in field studies (Honda et al., 2021b, 2021a; Qiu et al., 2017; Undap et al., 2013). However, the effects of individual pollutants must be assessed under controlled laboratory conditions to accurately evaluate the suitability of this species as an experimental model. This study introduces a novel approach for conducting laboratory experiments on wharf roaches. Further research should evaluate the impacts of various marine pollutants that accumulate in shoreline areas and the effects of long-term exposure. The wharf roach could serve as an excellent specimen for a coastal monitoring model to address the current knowledge gaps in this area.

Ingestion of microplastics by *Ligia* spp. may lead to changes in their behavior and symbiotic biota. In the present study, *Methanospirillum* and DNA viruses such as T4-like viruses and *Varicellovirus* were predicted as causal groups with optimal values. This observation is consistent with recent findings that viruses can modulate microbial communities in anaerobic digestion systems (Fan et al., 2023) and that diverse archaeal viruses can infect methanogens, including novel families such as *Speroviridae* and spindle-shaped viruses (Ngo et al., 2022). However, factor analysis showed that the value of h2, which indicates coordination in the other factor groups, was not low (h2 > 0.3; Table S3). As it was predicted that the detected factor group worked in cooperation with the calculation, the structural equation model was calculated using other factor groups as a starting point from the groove that formed the optimal structural equation model shown in Fig. 6b (Table S2). *Congregibacter*, a group of bacteria, and *Trichoplax*, a eukaryotic factor, were used to calculate the structural equations (Fig. S5), but the models composed of them show poor-suitability indices (Nos. 3 and 4, respectively) (Table S4). In addition, the number of successful bootstrap draws was low in this study.

Therefore, the relationship between the selected factors in Fig. S5 was quantitatively evaluated using BayesLiNGAM and a Bayesian score-based model. At least 56.71% or 28.47% of the relationships between the two viruses was calculated beyond the direct relationship between the two viruses (T4-like viruses and *Varicellovirus*), accounting for more than 80%, indicating that it is a causal group of high importance.

Furthermore, the notable increase in the abundance of *Methanospirillum*, a methane-producing archaeon, in the EPS group was highly intriguing. Methane, a primary driver of global warming, may also contribute to methane production through the consumption of materials such as EPS by wharf roaches in the environment, in addition to emissions originating from industrial settings and livestock facilities. However, whether the quantity of methane emitted by these wharf roaches is significant in relation to overall emissions remains unclear. Consequently, further experimentation is necessary to assess the methane production associated with wharf roach EPS ingestion in these animals. Nevertheless, the potential for methane generation following EPS ingestion suggests the utilization of EPS or certain additives as carbon sources, which warrants further investigation.

The present study confirmed that EPS ingestion induces transcriptional changes in the digestive tract and alters the gut microbiota of wharf roaches. In our previous studies, we revealed that wharf roaches are exposed to various chemical pollutants (Honda et al., 2021b, 2021a; Qiu et al., 2017; Undap et al., 2013) and that EPS ingestion frequently occurs in natural environments (Lee et al., 2025). Plastics are highly hydrophobic materials that can adsorb various pollutants (Batool and Valiyaveettil, 2021; Fu et al., 2021; Yu et al., 2019). In particular, EPS can adsorb a wider range of organic substances than other types of plastics owing to its porous structure and large surface area (Bhagwat et al., 2021; Li et al., 2020). In other words, the continuous exposure of wharf roaches to such complex pollutants in rocky coastal areas could exert more severe effects on their health. Moreover, organisms that act as decomposers typically play a crucial role in carbon cycling within the environment by ingesting and excreting organic matter. Therefore, changes in the gut microbiota can also affect other organisms and microorganisms in marine rocky habitats.

In conclusion, our study clearly revealed that wharf roaches consume polystyrene on rocky seashores and during feeding experiments. Dietary exposure test results confirmed that EPS ingestion caused gene expression changes and gut dysbiosis in wharf roaches, which transformed the microbiome flora and particularly increased the abundance of the methane producer *Methanospirillum*. These physiological and microbiological perturbations have the potential to propagate to ecosystem processes, underscoring the need for a systematic assessment of plastic-pollution risks across rocky-shore and coastal ecosystems.

## Data Availability

The sequencing data generated in this study, including RNA-seq of gut tissue and shotgun metagenome of gut microbiota from wharf roach (*Ligia sp.*), are available in the DDBJ Sequence Read Archive (DRA) under the BioProject accession number **PRJDB37655** (https://ddbj.nig.ac.jp/).

## Acknowledgements

This study was supported by Environment Research and Technology Development Fund [grant number JPMEERF20231001] and Grants-in-Aid for Scientific Research [JSPS KAKENHI Grant numbers JP21H05058 and JP22H03760].

We thank editors from Editage (https://www.editage.jp/), for English proofreading of the manuscript.

## Author contributions

**Seokhyun Lee**: Writing – original draft, Investigation, Methodology, Conceptualization, Formal analysis, Data curation

**Hirokuni Miyamoto**: Writing – review & editing, Writing – original draft, Formal analysis. **Yuki Takai**: Methodology, Formal analysis. **Wataru Suda** : Methodology, Writhing – original draft, **Hiroshi Ohno** : Methodology, **Yohei Simasaki**: Writing – review & editing, **Yuji Oshima**: Writing – review & editing, Supervision, Funding acquisition, Conceptualization

## Notes

### Competing Interest Statement

The authors have declared no competing interest.

## References

Assas, M., Qiu, X., Chen, K., Ogawa, H., Xu, H., Shimasaki, Y., Oshima, Y., 2020. Bioaccumulation and reproductive effects of fluorescent microplastics in medaka fish. Mar. Pollut. Bull. 158, 111446. 10.1016/j.marpolbul.2020.111446

Bajt, O., 2021. From plastics to microplastics and organisms. FEBS Open Bio. 11, 954–966. 10.1002/2211-5463.13120

Batool, A., Valiyaveettil, S., 2021. Surface functionalized cellulose fibers - A renewable adsorbent for removal of plastic nanoparticles from water. J. Hazard. Mater. 413, 125301. 10.1016/j.jhazmat.2021.125301

Bhagwat, G., Zhu, Q., O’Connor, W., Subashchandrabose, S., Grainge, I., Knight, R., Palanisami, T., 2021. Exploring the composition and functions of plastic microbiome using whole-genome sequencing. Environ. Sci. Technol. 55, 4899–4913. 10.1021/acs.est.0c07952

Bouchon, D., Zimmer, M., Dittmer, J., 2016. The terrestrial isopod microbiome: An all-in-one toolbox for animal-microbe interactions of ecological relevance. Front. Microbiol. 7, 1472. 10.3389/fmicb.2016.01472

Breiman, L., 2001. Random forests. Mach. Learn. 45, 5–32. 10.1023/a:1010933404324

Chen, Q., Yin, D., Jia, Y., Schiwy, S., Legradi, J., Yang, S., Hollert, H., 2017. Enhanced uptake of BPA in the presence of nanoplastics can lead to neurotoxic effects in adult zebrafish. Sci. Total Environ. 609, 1312–1321. 10.1016/j.scitotenv.2017.07.144

Choi, Y., Shin, D., Hong, C.P., Shin, D.-M., Cho, S.-H., Kim, S.S., Bae, M.A., Hong, S.H., Jang, M., Cho, Y., Han, G.M., Shim, W.J., Jung, J.-H., 2023. The effects of environmental microplastic on wharf roach (Ligia exotica): A multi-omics approach. Chemosphere 335, 139122. 10.1016/j.chemosphere.2023.139122

Fackelmann, G., Sommer, S., 2019. Microplastics and the gut microbiome: How chronically exposed species may suffer from gut dysbiosis. Mar. Pollut. Bull. 143, 193–203. 10.1016/j.marpolbul.2019.04.030

Fan, L., Peng, W., Duan, H., Lü, F., Zhang, H., He, P., 2023. Presence and role of viruses in anaerobic digestion of food waste under environmental variability. Microbiome 11, 170. 10.1186/s40168-023-01585-z

Fu, J., Zhou, S., Xu, H., Liao, L., Shen, H., Du, P., Zheng, X., 2023. ATM-ESCO2-SMC3 axis promotes 53BP1 recruitment in response to DNA damage and safeguards genome integrity by stabilizing cohesin complex. Nucleic Acids Res. 51, 7376–7391. 10.1093/nar/gkad533

Fu, L., Li, J., Wang, G., Luan, Y., Dai, W., 2021. Adsorption behavior of organic pollutants on microplastics. Ecotoxicol. Environ. Saf. 217, 112207. 10.1016/j.ecoenv.2021.112207

Gao, B., Shi, X., Li, S., Xu, W., Gao, N., Shan, J., Shen, W., 2023. Size-dependent effects of polystyrene microplastics on gut metagenome and antibiotic resistance in C57BL/6 mice. Ecotoxicol. Environ. Saf. 254, 114737. 10.1016/j.ecoenv.2023.114737

Heffner, T., Brami, S.A., Mendes, L.W., Kaupper, T., Hannula, E.S., Poehlein, A., Horn, M.A., Ho, A., 2023. Interkingdom interaction: the soil isopod Porcellio scaber stimulates the methane-driven bacterial and fungal interaction. ISME Commun. 3, 62. 10.1038/s43705-023-00271-3

Hink, L., Holzinger, A., Sandfeld, T., Weig, A.R., Schramm, A., Feldhaar, H., Horn, M.A., 2023. Microplastic ingestion affects hydrogen production and microbiomes in the gut of the terrestrial isopod Porcellio scaber. Environ. Microbiol. 25, 2776–2791. 10.1111/1462-2920.16386

Hodkovicova, N., Hollerova, A., Svobodova, Z., Faldyna, M., Faggio, C., 2022. Effects of plastic particles on aquatic invertebrates and fish - A review. Environ. Toxicol. Pharmacol. 96, 104013. 10.1016/j.etap.2022.104013

Honda, M., Mukai, K., Nagato, E., Uno, S., Oshima, Y., 2021a. Correlation between polycyclic aromatic hydrocarbons in wharf roach (Ligia spp.) and environmental components of the intertidal and supralittoral zone along the Japanese coast. Int. J. Environ. Res. Public Health 18, 630. 10.3390/ijerph18020630

Honda, M., Qiu, X., Lydia Undap, S., Kimura, T., Hori, T., Shimasaki, Y., Oshima, Y., 2021b. Contamination of heavy metals, polychlorinated dibenzo-p-dioxins/furans and dioxin-like polychlorinated biphenyls in wharf roach Ligia spp. In Japanese intertidal and supratidal zones. Appl. Sci. (Basel) 11, 1856. 10.3390/app11041856

Hoyer, P., Hyttinen, A., 2009. Bayesian discovery of linear acyclic causal models. In Proceedings of the Twenty-Fifth Conference on Uncertainty in Artificial Intelligence (UAI2009). 10.5555/1795114.1795143

Kazmi, S.S.U.H., Tayyab, M., Pastorino, P., Barcelò, D., Yaseen, Z.M., Grossart, H.-P., Khan, Z.H., Li, G., 2024. Decoding the molecular concerto: Toxicotranscriptomic evaluation of microplastic and nanoplastic impacts on aquatic organisms. J. Hazard. Mater. 472, 134574. 10.1016/j.jhazmat.2024.134574

Kuroda, M., Isobe, A., Uchida, K., Tokai, T., Kitakado, T., Yoshitake, M., Miyamoto, Y., Mukai, T., Imai, K., Shimizu, K., Yagi, M., Mituhasi, T., Habano, A., 2024. Abundance and potential sources of floating polystyrene foam macro- and microplastics around Japan. Sci. Total Environ. 925, 171421. 10.1016/j.scitotenv.2024.171421

Lee, S., Tsuruda, Y., Honda, M., Mukai, K., Hirasawa, T., Wijaya, D.C., Takai, Y., Simasaki, Y., Oshima, Y., 2025. Fragmentation of expanded polystyrene to microplastics by wharf roach Ligia spp. Mar. Pollut. Bull. 214, 117769. 10.1016/j.marpolbul.2025.117769

Lee, Y.-M., Choi, K.-M., Mun, S.H., Yoo, J.-W., Jung, J.-H., 2024. Gut microbiota composition of the isopod Ligia in South Korea exposed to expanded polystyrene pollution. PLoS One 19, e0308246. 10.1371/journal.pone.0308246

Li, M., Yu, H., Wang, Y., Li, J., Ma, G., Wei, X., 2020. QSPR models for predicting the adsorption capacity for microplastics of polyethylene, polypropylene and polystyrene. Sci. Rep. 10, 14597. 10.1038/s41598-020-71390-3

MacLeod, M., Arp, H.P.H., Tekman, M.B., Jahnke, A., 2021. The global threat from plastic pollution. Science 373, 61–65. 10.1126/science.abg5433

Medvedeva, S., Borrel, G., Krupovic, M., Gribaldo, S., 2023. A compendium of viruses from methanogenic archaea reveals their diversity and adaptations to the gut environment. Nat. Microbiol. 8, 2170–2182. 10.1038/s41564-023-01485-w

Miyamoto, H., Kikuchi, J., 2023. An evaluation of homeostatic plasticity for ecosystems using an analytical data science approach. Comput. Struct. Biotechnol. J. 21, 869–878. 10.1016/j.csbj.2023.01.001

Miyamoto, Hirokuni, Kawachi, N., Kurotani, A., Moriya, S., Suda, W., Suzuki, K., Matsuura, M., Tsuji, N., Nakaguma, T., Ishii, C., Tsuboi, A., Shindo, C., Kato, T., Udagawa, M., Satoh, T., Wada, S., Masuya, H., Miyamoto, Hisashi, Ohno, H., Kikuchi, J., 2023. Computational estimation of sediment symbiotic bacterial structures of seagrasses overgrowing downstream of onshore aquaculture. Environ. Res. 219, 115130. 10.1016/j.envres.2022.115130

Miyamoto, Hirokuni, Kawachi, N., Kurotani, A., Moriya, S., Suda, W., Suzuki, K., Matsuura, M., Tsuji, N., Nakaguma, T., Ishii, C., Tsuboi, A., Shindo, C., Kato, T., Udagawa, M., Satoh, T., Wada, S., Masuya, H., Miyamoto, Hisashi, Ohno, H., Kikuchi, J., 2022. Estimation of symbiotic bacterial structure in a sustainable seagrass ecosystem on recycled management. arXiv [q-bio.PE]. 10.1016/j.envres.2022.115130

Ngo, V.Q.H., Enault, F., Midoux, C., Mariadassou, M., Chapleur, O., Mazéas, L., Loux, V., Bouchez, T., Krupovic, M., Bize, A., 2022. Diversity of novel archaeal viruses infecting methanogens discovered through coupling of stable isotope probing and metagenomics. Environ. Microbiol. 24, 4853–4868. 10.1111/1462-2920.16120

Nishiyama, T., Ladurner, R., Schmitz, J., Kreidl, E., Schleiffer, A., Bhaskara, V., Bando, M., Shirahige, K., Hyman, A.A., Mechtler, K., Peters, J.-M., 2010. Sororin mediates sister chromatid cohesion by antagonizing Wapl. Cell 143, 737–749. 10.1016/j.cell.2010.10.031

Peprah, S., Ogwang, M.D., Kerchan, P., Reynolds, S.J., Tenge, C.N., Were, P.A., Kuremu, R.T., Wekesa, W.N., Masalu, N., Kawira, E., Otim, I., Legason, I.D., Ayers, L.W., Bhatia, K., Goedert, J.J., Pfeiffer, R.M., Mbulaiteye, S.M., 2021. Inverse association of falciparum positivity with endemic Burkitt lymphoma is robust in analyses adjusting for pre-enrollment malaria in the EMBLEM case-control study. Infect. Agent. Cancer 16, 40. 10.1186/s13027-021-00377-0

Qiu, X., Undap, S.L., Honda, M., Sekiguchi, T., Suzuki, N., Shimasaki, Y., Ando, H., Sato-Okoshi, W., Wada, T., Sunobe, T., Takeda, S., Munehara, H., Yokoyama, H., Momoshima, N., Oshima, Y., 2017. Pollution of radiocesium and radiosilver in wharf roach (Ligia sp.) by the Fukushima Dai-ichi Nuclear Power Plant accident. J. Radioanal. Nucl. Chem. 311, 121–126. 10.1007/s10967-016-4879-1

Quecholac-Piña, X., Hernández-Berriel, M.D.C., Mañón-Salas, M.D.C., Espinosa-Valdemar, R.M., Vázquez-Morillas, A., 2020. Degradation of plastics under anaerobic conditions: A short review. Polymers (Basel) 12, 109. 10.3390/polym12010109

Regoli, F., Giuliani, M.E., 2014. Oxidative pathways of chemical toxicity and oxidative stress biomarkers in marine organisms. Mar. Environ. Res. 93, 106–117. 10.1016/j.marenvres.2013.07.006

Revelle, W., Rocklin, T., 1979. Very Simple Structure: An alternative procedure for estimating the optimal number of interpretable factors. Multivariate Behav. Res. 14, 403–414. 10.1207/s15327906mbr1404_2

Rochman, C.M., Kurobe, T., Flores, I., Teh, S.J., 2014. Early warning signs of endocrine disruption in adult fish from the ingestion of polyethylene with and without sorbed chemical pollutants from the marine environment. Sci. Total Environ. 493, 656–661. 10.1016/j.scitotenv.2014.06.051

Rosseel, Y., 2012. Lavaan: An R package for structural equation modeling. J. Stat. Softw 48, 1–36. 10.18637/JSS.V048.I02

Rosseel, Y., Loh, W.W., 2022. A structural after measurement approach to structural equation modeling. Psychol. Methods. 10.1037/met0000503

Sagawa, N., Kawaai, K., Hinata, H., 2018. Abundance and size of microplastics in a coastal sea: Comparison among bottom sediment, beach sediment, and surface water. Mar. Pollut. Bull. 133, 532–542. 10.1016/j.marpolbul.2018.05.036

Schmitt, T.A., 2011. Current methodological considerations in exploratory and confirmatory factor analysis. J. Psychoeduc. Assess. 29, 304–321. 10.1177/0734282911406653

Shi, Z., Yao, F., Liu, Z., Zhang, J., 2024. Microplastics predominantly affect gut microbiota by altering community structure rather than richness and diversity: A meta-analysis of aquatic animals. Environ. Pollut. 360, 124639. 10.1016/j.envpol.2024.124639

Taguchi, Y., Kurotani, A., Yamano, H., Miyamoto, H., Kato, T., Tsuji, N., Matsuura, M., Nakaguma, T., Etoh, T., Shiotsuka, Y., Fujino, R., Udagawa, M., Kikuchi, J., Ohno, H., Takahashi, H., 2024. Causal estimation of maternal-offspring gut commensal bacterial associations under livestock grazing management conditions. Comput. Struct. Biotechnol. Rep. 1, 100012. 10.1016/j.csbr.2024.100012

Tamura, Y., Takai, Y., Miyamoto, H., SeokHyun, L., Liu, Y., Qiu, X., Kang, L.J., Simasaki, Y., Shindo, C., Suda, W., Ohno, H., Oshima, Y., 2024. Alteration of shoaling behavior and dysbiosis in the gut of medaka (Oryzias latipes) exposed to 2-μm polystyrene microplastics. Chemosphere 353, 141643. 10.1016/j.chemosphere.2024.141643

Tingley, D., Yamamoto, T., Hirose, K., Keele, L., Imai, K., 2014. Mediation:RPackage for causal mediation analysis. J. Stat. Softw. 59, 1–38. 10.18637/jss.v059.i05

Undap, S.L., Matsunaga, S., Honda, M., Sekiguchi, T., Suzuki, N., Khalil, F., Qiu, X., Shimasaki, Y., Ando, H., Sato-Okoshi, W., Sunobe, T., Takeda, S., Munehara, H., Oshima, Y., 2013. Accumulation of organotins in wharf roach (Ligia exotica Roux) and its ability to serve as a biomonitoring species for coastal pollution. Ecotoxicol. Environ. Saf. 96, 75–79. 10.1016/j.ecoenv.2013.06.019

Wang, K., Li, J., Zhao, L., Mu, X., Wang, C., Wang, M., Xue, X., Qi, S., Wu, L., 2021. Gut microbiota protects honey bees (Apis mellifera L.) against polystyrene microplastics exposure risks. J. Hazard. Mater. 402, 123828. 10.1016/j.jhazmat.2020.123828

Yang, Y., Kueh, A.J., Grant, Z.L., Abeysekera, W., Garnham, A.L., Wilcox, S., Hyland, C.D., Di Rago, L., Metcalf, D., Alexander, W.S., Coultas, L., Smyth, G.K., Voss, A.K., Thomas, T., 2022. The histone lysine acetyltransferase HBO1 (KAT7) regulates hematopoietic stem cell quiescence and self-renewal. Blood 139, 845–858. 10.1182/blood.2021013954

Yin, J., Liu, Z., Jin, X., Wang, W., Ma, L., Zhao, M., 2025. Characterization and regulatory analysis of betaine homocysteine S-methyltransferase gene 1 (BHMT1) in mud crab: A gene responsive to salinity and feeding behavior. Gene Rep. 38, 102102. 10.1016/j.genrep.2024.102102

Yu, F., Yang, C., Zhu, Z., Bai, X., Ma, J., 2019. Adsorption behavior of organic pollutants and metals on micro/nanoplastics in the aquatic environment. Sci. Total Environ. 694, 133643. 10.1016/j.scitotenv.2019.133643

Zhang, C., Li, F., Liu, X., Xie, L., Zhang, Y.T., Mu, J., 2023. Polylactic acid (PLA), polyethylene terephthalate (PET), and polystyrene (PS) microplastics differently affect the gut microbiota of marine medaka (Oryzias melastigma) after individual and combined exposure with sulfamethazine. Aquat. Toxicol. 259, 106522. 10.1016/j.aquatox.2023.106522

Zhirnov, O.P., Klenk, H.D., 2013. Influenza A virus proteins NS1 and hemagglutinin along with M2 are involved in stimulation of autophagy in infected cells. J. Virol. 87, 13107–13114. 10.1128/JVI.02148-13

Zhu, F., Li, D., Chen, K., 2019. Structures and functions of invertebrate glycosylation. Open Biol. 9, 180232. 10.1098/rsob.180232

## References

Bajt, O., 2021. From plastics to microplastics and organisms. FEBS Open Bio 11, 954–966. 10.1002/2211-5463.13120

Bhagwat, G., Zhu, Q., O’Connor, W., Subashchandrabose, S., Grainge, I., Knight, R., Palanisami, T., 2021. Exploring the Composition and Functions of Plastic Microbiome Using Whole-Genome Sequencing. Environ. Sci. Technol. 55, 4899–4913. 10.1021/acs.est.0c07952

Bouchon, D., Zimmer, M., Dittmer, J., 2016. The Terrestrial Isopod Microbiome: An All-in-One Toolbox for Animal-Microbe Interactions of Ecological Relevance. Front. Microbiol. 7, 1472. 10.3389/fmicb.2016.01472

Breiman, L., 2001. Random forests. Mach. Learn. 45, 5–32. 10.1023/a:1010933404324

Choi, Y., Shin, D., Hong, C.P., Shin, D.-M., Cho, S.-H., Kim, S.S., Bae, M.A., Hong, S.H., Jang, M., Cho, Y., Han, G.M., Shim, W.J., Jung, J.-H., 2023. The effects of environmental Microplastic on wharf roach (Ligia exotica): A Multi-Omics approach. Chemosphere 335, 139122. 10.1016/j.chemosphere.2023.139122

Hoyer, P., Hyttinen, A., 2009. Bayesian discovery of linear acyclic causal models. Uncertainty in Artificial Intelligence abs/1205.2641. 10.5555/1795114.1795143

Li, M., Yu, H., Wang, Y., Li, J., Ma, G., Wei, X., 2020. QSPR models for predicting the adsorption capacity for microplastics of polyethylene, polypropylene and polystyrene. Sci. Rep. 10. 10.1038/s41598-020-71390-3

Miyamoto, Hirokuni, Kawachi, N., Kurotani, A., Moriya, S., Suda, W., Suzuki, K., Matsuura, M., Tsuji, N., Nakaguma, T., Ishii, C., Tsuboi, A., Shindo, C., Kato, T., Udagawa, M., Satoh, T., Wada, S., Masuya, H., Miyamoto, Hisashi, Ohno, H., Kikuchi, J., 2022. Estimation of symbiotic bacterial structure in a sustainable seagrass ecosystem on recycled management. arXiv [q-bio.PE].

Rosseel, Y., 2012. Lavaan: An R package for structural equation modeling. Journal of Statistical Software 48, 1–36. 10.18637/JSS.V048.I02

Taguchi, Y., Kurotani, A., Yamano, H., Miyamoto, H., Kato, T., Tsuji, N., Matsuura, M., Nakaguma, T., Etoh, T., Shiotsuka, Y., Fujino, R., Udagawa, M., Kikuchi, J., Ohno, H., Takahashi, H., 2024. Causal estimation of maternal-offspring gut commensal bacterial associations under livestock grazing management conditions. Computational and Structural Biotechnology Reports 1, 100012. 10.1016/j.csbr.2024.100012

