## Supplement for "Changes in dysbiosis and gene expression in the gut of wharf roach (*Ligia* spp.) fed with expanded polystyrene": Supplementary.docx

Table S5.1 Gene expression changes induced by EPS exposure

| Seq-name | Description | logFC | P Value | FDR |
| --- | --- | --- | --- | --- |
| TRINITY_DN2890_c2_g1_i12 | Hemocyanin A chain [*Armadillidium vulgare*] | 12.963 | 1.82E-13 | 1.43E-09 |
| TRINITY_DN110213_c0_g1_i1 | Brachyurin [*Hyalella azteca*] | 11.973 | 1.19E-13 | 1.20E-09 |
| TRINITY_DN3241_c0_g1_i22 | Uncharacterized protein LOC123758464 isoform X3 [*Procambarus clarkii*] | 11.917 | 6.42E-12 | 2.64E-08 |
| TRINITY_DN1352_c0_g1_i2 | #N/A | 11.912 | 6.53E-12 | 2.64E-08 |
| TRINITY_DN352_c1_g1_i3 | #N/A | 11.383 | 3.94E-11 | 1.19E-07 |
| TRINITY_DN160_c0_g1_i4 | Hypothetical protein [*Idotea baltica*] | 11.328 | 4.74E-11 | 1.37E-07 |
| TRINITY_DN8597_c0_g1_i1 | #N/A | 11.093 | 1.05E-10 | 2.77E-07 |
| TRINITY_DN2297_c0_g2_i2 | Cytoplasmic dynein 1 intermediate chain 2-like isoform X26 [*Homarus americanus*] | 10.820 | 8.60E-12 | 3.25E-08 |
| TRINITY_DN9101_c0_g1_i5 | Extracellular matrix protein FRAS1 [*Armadillidium nasatum*] | 10.700 | 3.99E-10 | 8.62E-07 |
| TRINITY_DN16554_c0_g1_i20 | Echinoderm microtubule-associated protein-like 2 isoform X3 [*Penaeus vannamei*] | 10.630 | 5.08E-10 | 1.03E-06 |
| TRINITY_DN12783_c1_g1_i42 | General transcription factor II-I repeat domain-containing protein 2B-like [*Diabrotica virgifera virgifera*] | 10.230 | 1.95E-09 | 2.87E-06 |
| TRINITY_DN68950_c0_g1_i4 | Hypothetical protein [*Idotea baltica*] | 9.971 | 5.96E-12 | 2.64E-08 |
| TRINITY_DN3107_c0_g1_i5 | Autophagy-related protein 101-like [*Procambarus clarkii*] | 9.918 | 5.56E-09 | 7.15E-06 |
| TRINITY_DN4946_c0_g1_i8 | #N/A | 9.903 | 5.86E-09 | 7.39E-06 |
| TRINITY_DN7245_c0_g2_i1 | #N/A | 9.830 | 7.54E-09 | 8.94E-06 |
| TRINITY_DN5874_c0_g1_i2 | Zinc finger protein [*Armadillidium nasatum*] | 9.830 | 7.54E-09 | 8.94E-06 |
| TRINITY_DN28_c5_g3_i3 | #N/A | 9.799 | 8.34E-09 | 9.56E-06 |
| TRINITY_DN16534_c0_g1_i1 | #N/A | 9.566 | 1.81E-08 | 1.92E-05 |
| TRINITY_DN3893_c0_g1_i2 | Hypothetical protein Anas_07785, partial [*Armadillidium nasatum*] | 9.538 | 2.00E-08 | 2.05E-05 |
| TRINITY_DN1224_c0_g1_i40 | Influenza virus NS1A-binding protein homolog isoform X2 [*Cherax quadricarinatus*] | 9.513 | 2.15E-08 | 2.17E-05 |
| TRINITY_DN1236_c0_g1_i2 | Trypsin [*Hyalella azteca*] | 9.487 | 2.35E-08 | 2.33E-05 |
| TRINITY_DN807_c0_g1_i8 | DNA polymerase epsilon subunit 2 [*Nymphon striatum*] | 9.393 | 3.22E-08 | 3.00E-05 |
| TRINITY_DN120_c10_g1_i2 | #N/A | 9.106 | 8.33E-08 | 6.53E-05 |
| TRINITY_DN5963_c0_g1_i5 | NADH dehydrogenase [ubiquinone] 1 alpha subcomplex subunit 6 [*Schistocerca americana*] | 8.807 | 1.59E-07 | 1.07E-04 |
| TRINITY_DN10187_c0_g1_i4 | #N/A | 8.772 | 2.55E-07 | 1.59E-04 |
| TRINITY_DN4131_c0_g1_i10 | LOW-QUALITY PROTEIN: monocarboxylate transporter 14-like [*Penaeus monodon*] | 8.730 | 2.87E-07 | 1.77E-04 |
| TRINITY_DN12432_c0_g1_i36 | Calcium-transporting ATPase type 2C member 1, partial [*Armadillidium nasatum*] | 8.643 | 3.91E-07 | 2.23E-04 |
| TRINITY_DN5392_c0_g1_i11 | #N/A | 8.641 | 3.91E-07 | 2.23E-04 |
| TRINITY_DN15605_c0_g1_i6 | Zinc finger protein [*Armadillidium vulgare*] | 8.613 | 4.30E-07 | 2.37E-04 |
| TRINITY_DN1275_c0_g1_i8 | Mitotic checkpoint protein BUB3 [*Armadillidium vulgare*] | 8.612 | 4.30E-07 | 2.37E-04 |
| TRINITY_DN4893_c1_g1_i13 | Lysosomal-trafficking regulator-like isoform X1 [*Penaeus chinensis*] | 8.492 | 6.23E-07 | 3.19E-04 |
| TRINITY_DN10286_c0_g1_i4 | #N/A | 8.409 | 8.31E-07 | 4.06E-04 |
| TRINITY_DN9512_c1_g1_i2 | #N/A | 8.361 | 9.67E-07 | 4.64E-04 |
| TRINITY_DN101_c0_g1_i3 | Probable ATP-dependent RNA helicase DDX49 [*Homarus americanus*] | 8.359 | 9.67E-07 | 4.64E-04 |
| TRINITY_DN2635_c0_g1_i3 | A disintegrin and metalloproteinase with thrombospondin motifs 3-like [*Homarus americanus*] | 8.219 | 1.57E-06 | 6.86E-04 |
| TRINITY_DN15771_c0_g1_i6 | #N/A | 8.211 | 1.57E-06 | 6.86E-04 |
| TRINITY_DN9218_c1_g1_i1 | #N/A | 8.205 | 1.63E-06 | 7.01E-04 |
| TRINITY_DN44760_c0_g1_i1 | #N/A | 8.196 | 1.63E-06 | 7.01E-04 |
| TRINITY_DN772_c0_g1_i1 | #N/A | 8.030 | 2.80E-06 | 1.09E-03 |
| TRINITY_DN2748_c0_g1_i7 | Proton channel OtopLc-like isoform X8 [*Procambarus clarkii*] | 8.015 | 2.94E-06 | 1.13E-03 |
| TRINITY_DN57159_c0_g1_i2 | #N/A | 7.983 | 3.23E-06 | 1.21E-03 |
| TRINITY_DN2166_c0_g1_i42 | Hypothetical protein [*Idotea baltica*] | 7.842 | 5.13E-06 | 1.79E-03 |
| TRINITY_DN24362_c0_g1_i1 | #N/A | 7.744 | 1.68E-09 | 2.54E-06 |
| TRINITY_DN5153_c0_g2_i10 | Glutamate--cysteine ligase-like isoform X2 [*Homarus americanus*] | 7.724 | 1.86E-06 | 7.86E-04 |
| TRINITY_DN846_c0_g1_i10 | Palmitoyl-protein thioesterase 1 [*Portunus trituberculatus*] | 7.654 | 9.62E-06 | 2.98E-03 |
| TRINITY_DN367_c0_g1_i5 | CLIP-associating protein 2, partial [*Armadillidium nasatum*] | 7.650 | 9.62E-06 | 2.98E-03 |
| TRINITY_DN19882_c0_g1_i1 | Hypothetical protein [*Idotea baltica*] | 7.644 | 9.62E-06 | 2.98E-03 |
| TRINITY_DN36691_c0_g1_i11 | Dolichol kinase [*Armadillidium nasatum*] | 7.624 | 1.02E-05 | 3.08E-03 |
| TRINITY_DN2334_c1_g1_i10 | #N/A | 7.616 | 8.53E-08 | 6.53E-05 |
| TRINITY_DN630_c8_g1_i1 | #N/A | 7.613 | 1.09E-05 | 3.21E-03 |
| TRINITY_DN5662_c0_g1_i3 | #N/A | 7.601 | 1.09E-05 | 3.21E-03 |
| TRINITY_DN596_c9_g1_i2 | #N/A | 7.564 | 1.24E-05 | 3.54E-03 |
| TRINITY_DN71626_c0_g1_i3 | #N/A | 7.500 | 1.50E-05 | 4.06E-03 |
| TRINITY_DN7903_c1_g1_i1 | Uncharacterized protein LOC128702976 [*Cherax quadricarinatus*] | 7.477 | 1.61E-05 | 4.28E-03 |
| TRINITY_DN3961_c1_g3_i10 | #N/A | 7.471 | 1.61E-05 | 4.28E-03 |
| TRINITY_DN9248_c0_g1_i12 | Glycosyltransferase-like domain-containing protein 1-like [*Armadillidium nasatum*] | 7.463 | 1.72E-05 | 4.51E-03 |
| TRINITY_DN10800_c0_g1_i5 | #N/A | 7.425 | 1.98E-05 | 5.05E-03 |
| TRINITY_DN575_c0_g1_i2 | Hypothetical protein Anas_06632 [*Armadillidium nasatum*] | 7.322 | 4.17E-09 | 5.61E-06 |
| TRINITY_DN32587_c0_g1_i5 | Prolyl 4-hydroxylase subunit alpha-2, partial [*Armadillidium nasatum*] | 7.297 | 2.85E-05 | 6.83E-03 |
| TRINITY_DN3960_c0_g1_i3 | Hypothetical protein [*Idotea baltica*] | 7.286 | 2.85E-05 | 6.83E-03 |
| TRINITY_DN1355_c0_g1_i3 | #N/A | 7.285 | 2.85E-05 | 6.83E-03 |
| TRINITY_DN13610_c0_g2_i6 | Hypothetical protein [*Idotea baltica*] | 7.278 | 3.08E-05 | 7.31E-03 |
| TRINITY_DN9101_c0_g1_i2 | Extracellular matrix protein FRAS1 [*Armadillidium nasatum*] | 7.161 | 1.54E-07 | 1.05E-04 |
| TRINITY_DN65071_c0_g2_i1 | #N/A | 7.143 | 4.60E-05 | 1.02E-02 |
| TRINITY_DN74968_c0_g1_i2 | L-asparaginase [*Armadillidium vulgare*] | 7.132 | 4.60E-05 | 1.02E-02 |
| TRINITY_DN3556_c1_g1_i2 | Cathepsin C 1 [*Panulirus homarus*] | 7.124 | 9.74E-09 | 1.09E-05 |
| TRINITY_DN23828_c0_g1_i9 | Tectonin beta-propeller repeat-containing protein [*Armadillidium vulgare*] | 7.093 | 5.46E-05 | 1.13E-02 |
| TRINITY_DN24806_c0_g1_i6 | Protein transport protein Sec31B [*Armadillidium nasatum*] | 7.074 | 5.46E-05 | 1.13E-02 |
| TRINITY_DN6174_c0_g1_i4 | #N/A | 7.069 | 5.96E-05 | 1.22E-02 |
| TRINITY_DN1040_c0_g1_i10 | Copper-transporting ATPase 1 [*Armadillidium nasatum*] | 7.045 | 1.33E-06 | 6.17E-04 |
| TRINITY_DN2059_c0_g1_i8 | 3-Hydroxyanthranilate 3,4-dioxygenase-like [*Penaeus japonicus*] | 7.020 | 6.51E-05 | 1.32E-02 |
| TRINITY_DN11212_c1_g1_i2 | #N/A | 6.962 | 7.82E-05 | 1.51E-02 |
| TRINITY_DN3513_c0_g1_i6 | Protein max-like isoform X4 [*Homarus americanus*] | 6.958 | 7.82E-05 | 1.51E-02 |
| TRINITY_DN4056_c0_g1_i7 | Uncharacterized protein LOC126997656 [*Eriocheir sinensis*] | 6.844 | 6.78E-06 | 2.28E-03 |
| TRINITY_DN38284_c1_g2_i1 | #N/A | 6.818 | 1.28E-04 | 2.15E-02 |
| TRINITY_DN38722_c1_g3_i4 | Uncharacterized protein Anas_02879 [*Armadillidium nasatum*] | 6.668 | 1.96E-04 | 3.02E-02 |
| TRINITY_DN3119_c0_g4_i4 | #N/A | 6.613 | 2.20E-04 | 3.32E-02 |
| TRINITY_DN36465_c0_g1_i1 | V-type proton ATPase 16 kDa proteolipid subunit [*Armadillidium vulgare*] | 6.460 | 8.94E-08 | 6.76E-05 |
| TRINITY_DN3559_c0_g3_i4 | Putative 4-coumarate--CoA ligase-like 8 [*Armadillidium nasatum*] | 6.349 | 2.18E-07 | 1.40E-04 |
| TRINITY_DN7275_c3_g1_i2 | #N/A | 6.301 | 3.32E-06 | 1.23E-03 |
| TRINITY_DN399_c1_g1_i4 | Hypothetical protein [*Idotea baltica*] | 6.300 | 1.25E-07 | 9.14E-05 |
| TRINITY_DN832_c0_g1_i7 | #N/A | 6.261 | 3.26E-05 | 7.65E-03 |
| TRINITY_DN9_c0_g1_i19 | Salivary glue protein Sgs-3-like isoform X1 [*Cherax quadricarinatus*] | 6.259 | 1.32E-07 | 9.42E-05 |
| TRINITY_DN797_c0_g1_i10 | Hypothetical protein Anas_14230, partial [*Armadillidium nasatum*] | 5.952 | 3.96E-07 | 2.24E-04 |
| TRINITY_DN6052_c1_g1_i3 | Uncharacterized protein LOC118732360 [*Rhagoletis pomonella*] | 5.871 | 2.44E-06 | 9.80E-04 |
| TRINITY_DN25230_c0_g1_i1 | Ileal sodium/bile acid cotransporter [*Armadillidium nasatum*] | 5.725 | 2.39E-06 | 9.69E-04 |
| TRINITY_DN24584_c0_g1_i6 | Putative pyruvate dehydrogenase E1 component subunit alpha, mitochondrial [*Armadillidium nasatum*] | 5.689 | 1.80E-06 | 7.68E-04 |
| TRINITY_DN2353_c6_g1_i16 | #N/A | 5.665 | 6.83E-06 | 2.28E-03 |
| TRINITY_DN1833_c0_g1_i6 | Phospholipase ABHD3-like isoform X2 [*Portunus trituberculatus*] | 5.600 | 3.44E-05 | 7.98E-03 |
| TRINITY_DN474_c0_g1_i5 | Uncharacterized protein LOC122242129 [*Penaeus japonicus*] | 5.572 | 2.82E-06 | 1.10E-03 |
| TRINITY_DN89339_c1_g2_i10 | #N/A | 5.537 | 7.95E-05 | 1.52E-02 |
| TRINITY_DN3947_c1_g1_i1 | #N/A | 5.514 | 1.20E-04 | 2.05E-02 |
| TRINITY_DN53917_c3_g1_i8 | Beta-1,3-glucanase, partial [*Birgus latro*] | 5.381 | 2.47E-06 | 9.85E-04 |
| TRINITY_DN105608_c0_g1_i1 | Cytochrome oxidase subunit I, partial [*Aegla franciscana*] | 5.330 | 2.63E-06 | 1.04E-03 |
| TRINITY_DN2798_c0_g1_i1 | #N/A | 5.321 | 3.72E-06 | 1.35E-03 |
| TRINITY_DN1_c0_g2_i7 | Protein white-like isoform X3 [*Eriocheir sinensis*] | 5.265 | 1.02E-05 | 3.08E-03 |
| TRINITY_DN4038_c0_g1_i1 | Hypothetical protein Avbf_00356 [*Armadillidium vulgare*] | 5.216 | 4.77E-06 | 1.69E-03 |
| TRINITY_DN1957_c0_g2_i19 | Sestrin-2-like isoform X3 [*Homarus americanus*] | 5.202 | 3.96E-06 | 1.42E-03 |
| TRINITY_DN3525_c0_g1_i7 | N-acetylated-alpha-linked acidic dipeptidase-like protein [*Armadillidium nasatum*] | 5.165 | 4.85E-05 | 1.04E-02 |
| TRINITY_DN4946_c0_g1_i7 | #N/A | 5.144 | 8.18E-06 | 2.70E-03 |
| TRINITY_DN4750_c0_g1_i23 | Hypothetical protein Avbf_00926 [*Armadillidium vulgare*] | 5.080 | 8.37E-05 | 1.58E-02 |
| TRINITY_DN1628_c0_g2_i6 | Monocarboxylate transporter 12, partial [*Armadillidium nasatum*] | 5.065 | 9.80E-06 | 3.02E-03 |
| TRINITY_DN5226_c0_g1_i3 | Thioredoxin reductase 3 [*Armadillidium nasatum*] | 5.055 | 5.89E-06 | 2.05E-03 |
| TRINITY_DN8019_c0_g1_i3 | Pancreatic triacylglycerol lipase-like [*Homarus americanus*] | 5.048 | 2.02E-05 | 5.15E-03 |
| TRINITY_DN388_c1_g1_i2 | #N/A | 5.038 | 2.81E-04 | 4.04E-02 |
| TRINITY_DN3670_c3_g1_i4 | #N/A | 5.007 | 4.51E-05 | 1.02E-02 |
| TRINITY_DN75649_c0_g1_i1 | #N/A | 4.971 | 1.69E-05 | 4.46E-03 |
| TRINITY_DN474_c0_g1_i12 | Uncharacterized protein LOC125029819 [*Penaeus chinensis*] | 4.968 | 1.16E-05 | 3.35E-03 |
| TRINITY_DN23578_c0_g1_i1 | #N/A | 4.956 | 7.89E-05 | 1.52E-02 |
| TRINITY_DN3973_c0_g1_i20 | Tensin-1, partial [*Armadillidium vulgare*] | 4.837 | 2.21E-05 | 5.54E-03 |
| TRINITY_DN8525_c0_g2_i11 | Iodotyrosine deiodinase-like isoform X3 [*Eriocheir sinensis*] | 4.837 | 4.77E-05 | 1.03E-02 |
| TRINITY_DN2673_c0_g1_i4 | Solute carrier family 13 member 5-like isoform X2 [*Penaeus monodon*] | 4.830 | 1.37E-05 | 3.92E-03 |
| TRINITY_DN2833_c0_g1_i5 | #N/A | 4.828 | 1.23E-04 | 2.08E-02 |
| TRINITY_DN1822_c0_g2_i9 | #N/A | 4.811 | 1.50E-05 | 4.06E-03 |
| TRINITY_DN3308_c0_g1_i2 | #N/A | 4.806 | 2.08E-04 | 3.17E-02 |
| TRINITY_DN2940_c0_g1_i8 | Hypothetical protein C7M84_014775 [*Penaeus vannamei*] | 4.795 | 2.52E-05 | 6.20E-03 |
| TRINITY_DN34360_c0_g2_i3 | Zinc finger CCCH domain-containing protein 18 [*Armadillidium vulgare*] | 4.764 | 2.06E-05 | 5.22E-03 |
| TRINITY_DN2259_c0_g1_i4 | Hypothetical protein Avbf_02172 [*Armadillidium vulgare*] | 4.753 | 2.18E-05 | 5.49E-03 |
| TRINITY_DN4368_c0_g1_i12 | Betaine--homocysteine S-methyltransferase 1-like [*Penaeus monodon*] | 4.749 | 1.55E-05 | 4.16E-03 |
| TRINITY_DN1556_c0_g1_i14 | #N/A | 4.747 | 2.30E-05 | 5.74E-03 |
| TRINITY_DN39263_c0_g1_i8 | Rhomboid-related protein 4 [*Armadillidium nasatum*] | 4.707 | 6.62E-05 | 1.33E-02 |
| TRINITY_DN4139_c0_g1_i6 | Cytosolic phospholipase A2 [*Armadillidium vulgare*] | 4.684 | 2.26E-05 | 5.64E-03 |
| TRINITY_DN664_c0_g1_i1 | XP_045603483.1 uncharacterized protein LOC123761500 [*Procambarus clarkii*] | 4.675 | 4.59E-05 | 1.02E-02 |
| TRINITY_DN492_c1_g1_i4 | #N/A | 4.653 | 2.68E-05 | 6.51E-03 |
| TRINITY_DN1_c0_g2_i15 | Hypothetical protein [*Idotea baltica*] | 4.621 | 8.98E-05 | 1.64E-02 |
| TRINITY_DN5503_c0_g1_i7 | CAAX prenyl protease 1-like protein, partial [*Armadillidium vulgare*] | 4.618 | 4.48E-05 | 1.02E-02 |
| TRINITY_DN29814_c0_g2_i6 | #N/A | 4.616 | 5.70E-05 | 1.17E-02 |
| TRINITY_DN4888_c0_g1_i17 | #N/A | 4.584 | 6.60E-05 | 1.33E-02 |
| TRINITY_DN68926_c0_g1_i13 | Cytochrome P450 302a1, mitochondrial-like [*Penaeus monodon*] | 4.574 | 4.77E-05 | 1.03E-02 |
| TRINITY_DN3065_c1_g1_i6 | Period [*Eurydice pulchra*] | 4.565 | 3.14E-05 | 7.41E-03 |
| TRINITY_DN2061_c0_g5_i1 | Basigin [*Armadillidium nasatum*] | 4.521 | 5.66E-05 | 1.17E-02 |
| TRINITY_DN91453_c0_g1_i1 | Hypothetical protein [*Idotea baltica*] | 4.513 | 3.54E-05 | 8.17E-03 |
| TRINITY_DN8618_c0_g3_i1 | Ubiquitin carboxyl-terminal hydrolase 8 [*Armadillidium vulgare*] | 4.501 | 7.24E-05 | 1.43E-02 |
| TRINITY_DN3289_c0_g1_i20 | #N/A | 4.494 | 3.73E-05 | 8.57E-03 |
| TRINITY_DN1680_c0_g1_i2 | #N/A | 4.493 | 7.60E-05 | 1.49E-02 |
| TRINITY_DN5428_c0_g1_i5 | Hypothetical protein Avbf_06013 [*Armadillidium vulgare*] | 4.492 | 4.75E-05 | 1.03E-02 |
| TRINITY_DN5528_c0_g1_i3 | Carboxypeptidase activation peptide [*Trinorchestia longiramus*] | 4.488 | 4.78E-05 | 1.03E-02 |
| TRINITY_DN148_c0_g1_i116 | Basement membrane-specific heparan Sulfate proteoglycan core protein, partial [*Armadillidium nasatum*] | 4.481 | 5.09E-05 | 1.07E-02 |
| TRINITY_DN946_c0_g1_i13 | Myeloid leukemia factor-like [*Penaeus japonicus*] | 4.465 | 7.16E-05 | 1.42E-02 |
| TRINITY_DN2373_c0_g1_i5 | Ephrin type-B receptor 1-B-like isoform X8 [*Penaeus japonicus*] | 4.458 | 1.01E-04 | 1.80E-02 |
| TRINITY_DN9218_c0_g2_i2 | Venom phosphodiesterase 2 [*Armadillidium nasatum*] | 4.430 | 4.65E-05 | 1.02E-02 |
| TRINITY_DN531_c2_g4_i2 | Hypothetical protein Anas_07814 [*Armadillidium nasatum*] | 4.410 | 4.97E-05 | 1.06E-02 |
| TRINITY_DN6995_c0_g1_i16 | Uncharacterized protein LOC108680125 [*Hyalella azteca*] | 4.407 | 9.40E-05 | 1.71E-02 |
| TRINITY_DN6023_c0_g1_i57 | Transmembrane protein 11, mitochondrial-like [*Penaeus monodon*] | 4.387 | 3.13E-04 | 4.37E-02 |
| TRINITY_DN110_c0_g1_i3 | 2-Acylglycerol O-acyltransferase 1-like [*Eriocheir sinensis*] | 4.381 | 3.24E-04 | 4.50E-02 |
| TRINITY_DN5477_c0_g1_i9 | LOW-QUALITY PROTEIN: NFX1-Type zinc finger-containing protein 1-like [*Cherax quadricarinatus*] | 4.375 | 7.75E-05 | 1.51E-02 |
| TRINITY_DN5662_c0_g2_i1 | #N/A | 4.341 | 1.66E-04 | 2.64E-02 |
| TRINITY_DN420_c1_g1_i1 | Cytochrome P450 [*Armadillidium nasatum*] | 4.315 | 8.61E-05 | 1.61E-02 |
| TRINITY_DN2569_c0_g1_i3 | #N/A | 4.304 | 6.57E-05 | 1.33E-02 |
| TRINITY_DN1328_c0_g1_i4 | G patch domain-containing protein 2-like [*Procambarus clarkii*] | 4.299 | 1.04E-04 | 1.84E-02 |
| TRINITY_DN223_c0_g1_i88 | #N/A | 4.289 | 1.10E-04 | 1.92E-02 |
| TRINITY_DN32130_c0_g1_i21 | Sorbitol dehydrogenase-like [*Penaeus japonicus*] | 4.273 | 7.38E-05 | 1.45E-02 |
| TRINITY_DN3099_c0_g1_i8 | E3 ubiquitin-protein ligase UBR4-like isoform X3 [*Homarus americanus*] | 4.255 | 8.69E-05 | 1.61E-02 |
| TRINITY_DN8273_c0_g1_i9 | Dehydrogenase/reductase SDR family member 11 [*Armadillidium vulgare*] | 4.255 | 1.21E-04 | 2.07E-02 |
| TRINITY_DN4274_c0_g1_i7 | Methionine synthase reductase [*Armadillidium nasatum*] | 4.238 | 1.90E-04 | 2.94E-02 |
| TRINITY_DN797_c0_g1_i12 | Hypothetical protein Anas_14230, partial [*Armadillidium nasatum*] | 4.202 | 9.88E-05 | 1.77E-02 |
| TRINITY_DN24578_c0_g1_i3 | DNA damage-regulated autophagy modulator protein 2 [*Armadillidium nasatum*] | 4.187 | 2.68E-04 | 3.92E-02 |
| TRINITY_DN1297_c0_g1_i8 | Chondroitin sulfate synthase 1-like [*Penaeus chinensis*] | 4.184 | 1.60E-04 | 2.59E-02 |
| TRINITY_DN1315_c1_g1_i1 | #N/A | 4.182 | 1.84E-04 | 2.87E-02 |
| TRINITY_DN11543_c0_g1_i19 | Hypothetical protein [*Idotea baltica*] | 4.176 | 2.34E-04 | 3.49E-02 |
| TRINITY_DN271_c0_g1_i17 | Phosphatidylinositol 4-phosphate 5-kinase type-1 gamma-like isoform X9 [*Procambarus clarkii*] | 4.176 | 1.75E-04 | 2.78E-02 |
| TRINITY_DN2652_c0_g1_i15 | #N/A | 4.166 | 1.65E-04 | 2.64E-02 |
| TRINITY_DN4552_c1_g2_i10 | #N/A | 4.152 | 2.12E-04 | 3.22E-02 |
| TRINITY_DN8030_c2_g1_i7 | RNA-directed DNA polymerase from mobile element jockey-like protein [*Dinothrombium tinctorium*] | 4.132 | 1.87E-04 | 2.91E-02 |
| TRINITY_DN1237_c2_g1_i5 | Hypothetical protein [*Idotea baltica*] | 4.126 | 1.10E-04 | 1.92E-02 |
| TRINITY_DN3183_c2_g1_i1 | #N/A | 4.120 | 1.51E-04 | 2.46E-02 |
| TRINITY_DN4522_c0_g1_i27 | Acetyl-CoA carboxylase [*Armadillidium nasatum*] | 4.097 | 1.85E-04 | 2.88E-02 |
| TRINITY_DN6192_c0_g1_i2 | Serine/threonine-protein kinase pim-1 [*Armadillidium nasatum*] | 4.079 | 1.50E-04 | 2.46E-02 |
| TRINITY_DN16534_c0_g1_i3 | #N/A | 4.078 | 1.49E-04 | 2.45E-02 |
| TRINITY_DN704_c0_g2_i1 | Hypothetical protein B7P43_G11307 [*Cryptotermes secundus*] | 4.077 | 1.77E-04 | 2.79E-02 |
| TRINITY_DN60692_c0_g1_i1 | #N/A | 4.027 | 3.48E-04 | 4.75E-02 |
| TRINITY_DN58119_c0_g1_i2 | Elongation factor 1-alpha 1 [*Armadillidium vulgare*] | 4.011 | 1.64E-04 | 2.63E-02 |
| TRINITY_DN1233_c0_g1_i1 | Importin subunit alpha-3 [*Armadillidium vulgare*] | 3.977 | 2.80E-04 | 4.03E-02 |
| TRINITY_DN2525_c0_g2_i5 | Endoglucanase, partial [*Porcellio scaber*] | 3.960 | 3.21E-04 | 4.46E-02 |
| TRINITY_DN759_c1_g1_i21 | Serine proteases trypsin domain, partial [*Trinorchestia longiramus*] | 3.951 | 3.20E-04 | 4.46E-02 |
| TRINITY_DN213_c1_g1_i10 | Endoglucanase 1 [*Armadillidium nasatum*] | 3.937 | 1.99E-04 | 3.05E-02 |
| TRINITY_DN27765_c0_g1_i10 | Procathepsin L-like [*Eriocheir sinensis*] | 3.927 | 3.63E-04 | 4.91E-02 |
| TRINITY_DN508_c0_g1_i5 | Hypothetical protein Avbf_00356 [*Armadillidium vulgare*] | 3.901 | 2.66E-04 | 3.90E-02 |
| TRINITY_DN13315_c0_g2_i2 | Uncharacterized protein LOC128984270 [*Macrosteles quadrilineatus*] | 3.889 | 3.05E-04 | 4.28E-02 |
| TRINITY_DN1831_c5_g1_i3 | Ester hydrolase C11orf54-like protein [*Armadillidium vulgare*] | 3.828 | 2.78E-04 | 4.03E-02 |
| TRINITY_DN1408_c0_g1_i1 | Pyridoxine-5'-phosphate oxidase [*Armadillidium vulgare*] | 3.817 | 2.98E-04 | 4.19E-02 |
| TRINITY_DN5528_c0_g3_i2 | Carboxypeptidase activation peptide [*Trinorchestia longiramus*] | 3.784 | 3.34E-04 | 4.59E-02 |
| TRINITY_DN2403_c0_g1_i1 | #N/A | -3.756 | 3.71E-04 | 5.00E-02 |
| TRINITY_DN3647_c2_g1_i1 | #N/A | -3.757 | 3.55E-04 | 4.82E-02 |
| TRINITY_DN7400_c0_g1_i7 | #N/A | -3.780 | 3.56E-04 | 4.83E-02 |
| TRINITY_DN9092_c0_g1_i27 | Hypothetical protein HF086_004231 [*Spodoptera exigua*] | -3.802 | 3.55E-04 | 4.82E-02 |
| TRINITY_DN81510_c2_g1_i1 | Betaine--homocysteine S-methyltransferase 1-like [*Penaeus monodon*] | -3.832 | 2.78E-04 | 4.03E-02 |
| TRINITY_DN12873_c0_g1_i1 | #N/A | -3.837 | 2.97E-04 | 4.19E-02 |
| TRINITY_DN956_c0_g1_i6 | Hypothetical protein Anas_02118 [*Armadillidium nasatum*] | -3.844 | 3.33E-04 | 4.59E-02 |
| TRINITY_DN17329_c1_g1_i3 | #N/A | -3.859 | 2.84E-04 | 4.06E-02 |
| TRINITY_DN20103_c0_g1_i4 | Probable chitinase 10 [*Hyalella azteca*] | -3.862 | 2.97E-04 | 4.19E-02 |
| TRINITY_DN2076_c0_g3_i7 | #N/A | -3.872 | 3.28E-04 | 4.54E-02 |
| TRINITY_DN2202_c0_g1_i3 | Hypothetical protein Avbf_05560 [*Armadillidium vulgare*] | -3.880 | 3.43E-04 | 4.70E-02 |
| TRINITY_DN1561_c0_g1_i25 | Meiosis regulator and mRNA stability factor 1-like isoform X3 [*Procambarus clarkii*] | -3.892 | 2.80E-04 | 4.03E-02 |
| TRINITY_DN24559_c0_g1_i1 | Gametocyte-specific factor 1, partial [*Armadillidium vulgare*] | -3.899 | 2.30E-04 | 3.45E-02 |
| TRINITY_DN9028_c0_g1_i5 | Hypothetical protein Avbf_13578 [*Armadillidium vulgare*] | -3.919 | 3.64E-04 | 4.91E-02 |
| TRINITY_DN7245_c0_g2_i3 | #N/A | -3.925 | 3.29E-04 | 4.55E-02 |
| TRINITY_DN36354_c0_g1_i1 | #N/A | -3.932 | 2.86E-04 | 4.07E-02 |
| TRINITY_DN5064_c0_g1_i1 | Low-density lipoprotein receptor [*Armadillidium nasatum*] | -3.948 | 2.21E-04 | 3.33E-02 |
| TRINITY_DN697_c0_g1_i24 | Liprin-alpha-2 [*Armadillidium nasatum*] | -3.960 | 2.52E-04 | 3.73E-02 |
| TRINITY_DN5912_c0_g3_i2 | Protein rhomboid [*Armadillidium nasatum*] | -3.970 | 2.84E-04 | 4.06E-02 |
| TRINITY_DN48185_c2_g1_i3 | #N/A | -3.977 | 2.60E-04 | 3.83E-02 |
| TRINITY_DN586_c0_g2_i1 | #N/A | -3.988 | 2.41E-04 | 3.58E-02 |
| TRINITY_DN26822_c0_g4_i1 | Serine proteinase stubble-like isoform X3 [*Procambarus clarkii*] | -3.989 | 2.42E-04 | 3.59E-02 |
| TRINITY_DN2038_c0_g1_i16 | #N/A | -3.990 | 1.82E-04 | 2.85E-02 |
| TRINITY_DN2296_c0_g1_i4 | PAS domain-containing serine/threonine-protein kinase [*Armadillidium nasatum*] | -3.992 | 3.39E-04 | 4.65E-02 |
| TRINITY_DN1495_c10_g1_i5 | #N/A | -4.001 | 1.72E-04 | 2.74E-02 |
| TRINITY_DN1257_c1_g1_i11 | T-related protein [*Pseudolycoriella hygida*] | -4.013 | 1.64E-04 | 2.63E-02 |
| TRINITY_DN1850_c0_g1_i1 | #N/A | -4.015 | 2.85E-04 | 4.06E-02 |
| TRINITY_DN895_c0_g1_i8 | Erythroid differentiation-related factor 1 [*Armadillidium nasatum*] | -4.022 | 2.36E-04 | 3.52E-02 |
| TRINITY_DN34848_c0_g1_i1 | Hypothetical protein Avbf_10105 [*Armadillidium vulgare*] | -4.043 | 2.54E-04 | 3.74E-02 |
| TRINITY_DN14751_c1_g1_i44 | Tyrosine-protein phosphatase non-receptor type 14-like [*Penaeus monodon*] | -4.044 | 3.12E-04 | 4.36E-02 |
| TRINITY_DN11038_c0_g3_i12 | Twitchin-like isoform X14 [*Procambarus clarkii*] | -4.046 | 1.64E-04 | 2.63E-02 |
| TRINITY_DN1049_c0_g1_i2 | Lysosomal acid phosphatase, partial [*Armadillidium vulgare*] | -4.063 | 1.64E-04 | 2.63E-02 |
| TRINITY_DN57547_c0_g1_i2 | Protein Skeletor, isoforms B/C-like [*Penaeus japonicus*] | -4.089 | 2.61E-04 | 3.84E-02 |
| TRINITY_DN174181_c0_g1_i1 | Facilitated trehalose transporter Tret1-like isoform X2 [*Plodia interpunctella*] | -4.099 | 1.36E-04 | 2.27E-02 |
| TRINITY_DN2191_c26_g1_i1 | #N/A | -4.102 | 2.30E-04 | 3.45E-02 |
| TRINITY_DN3087_c1_g1_i4 | Hypothetical protein Anas_00004, partial [*Armadillidium nasatum*] | -4.148 | 1.41E-04 | 2.35E-02 |
| TRINITY_DN2869_c1_g4_i3 | Cryptochrome 2 [*Eurydice pulchra*] | -4.168 | 1.24E-04 | 2.10E-02 |
| TRINITY_DN1495_c10_g1_i2 | #N/A | -4.174 | 1.23E-04 | 2.08E-02 |
| TRINITY_DN2849_c0_g1_i3 | Hypothetical protein Anas_06632 [*Armadillidium nasatum*] | -4.175 | 9.52E-05 | 1.72E-02 |
| TRINITY_DN4946_c0_g1_i5 | #N/A | -4.182 | 1.19E-04 | 2.05E-02 |
| TRINITY_DN2297_c0_g3_i4 | Cytoplasmic dynein 1 intermediate chain 2-like isoform X26 [*Homarus americanus*] | -4.191 | 1.05E-04 | 1.85E-02 |
| TRINITY_DN88238_c0_g2_i1 | Uncharacterized protein LOC123769984 [*Procambarus clarkii*] | -4.195 | 2.18E-04 | 3.29E-02 |
| TRINITY_DN47_c0_g2_i10 | #N/A | -4.203 | 1.02E-04 | 1.81E-02 |
| TRINITY_DN457_c0_g1_i21 | #N/A | -4.206 | 9.68E-05 | 1.74E-02 |
| TRINITY_DN25444_c0_g1_i10 | Probable phosphorylase b kinase regulatory subunit alpha isoform X12 [*Homarus americanus*] | -4.216 | 1.59E-04 | 2.58E-02 |
| TRINITY_DN4135_c0_g1_i2 | Death-associated protein kinase related [*Armadillidium vulgare*] | -4.219 | 1.06E-04 | 1.85E-02 |
| TRINITY_DN2297_c0_g1_i3 | Protein phosphatase 1 regulatory subunit 3C-B [*Armadillidium vulgare*] | -4.221 | 1.14E-04 | 1.98E-02 |
| TRINITY_DN10509_c0_g5_i1 | #N/A | -4.222 | 1.82E-04 | 2.85E-02 |
| TRINITY_DN1937_c0_g3_i5 | Hypothetical protein [*Idotea baltica*] | -4.247 | 7.86E-05 | 1.51E-02 |
| TRINITY_DN576_c0_g1_i10 | Hypothetical protein Anas_07065 [*Armadillidium nasatum*] | -4.252 | 1.48E-04 | 2.44E-02 |
| TRINITY_DN2775_c5_g1_i1 | #N/A | -4.254 | 1.99E-04 | 3.05E-02 |
| TRINITY_DN4705_c0_g1_i7 | Tyrosine-protein kinase SRK2-like isoform X2 [*Eriocheir sinensis*] | -4.257 | 2.88E-04 | 4.10E-02 |
| TRINITY_DN18632_c0_g1_i2 | Venom protease-like, partial [*Daktulosphaira vitifoliae*] | -4.261 | 1.28E-04 | 2.15E-02 |
| TRINITY_DN777_c1_g1_i8 | Condensin-2 complex subunit G2-like [*Cherax quadricarinatus*] | -4.288 | 1.21E-04 | 2.07E-02 |
| TRINITY_DN5202_c0_g1_i1 | Rho GTPase-activating protein 20 [*Armadillidium vulgare*] | -4.298 | 7.33E-05 | 1.44E-02 |
| TRINITY_DN12448_c0_g1_i1 | Lysophospholipid acyltransferase 7 [*Armadillidium nasatum*] | -4.303 | 1.78E-04 | 2.80E-02 |
| TRINITY_DN113_c0_g2_i4 | E3 ubiquitin-protein ligase RBBP6-like, partial [*Penaeus monodon*] | -4.305 | 8.37E-05 | 1.58E-02 |
| TRINITY_DN17114_c0_g1_i1 | Caveolin-3-like isoform X3 [*Penaeus monodon*] | -4.356 | 9.43E-05 | 1.71E-02 |
| TRINITY_DN6370_c1_g2_i3 | Multivesicular body subunit 12B [*Armadillidium vulgare*] | -4.370 | 2.00E-04 | 3.05E-02 |
| TRINITY_DN1899_c0_g1_i5 | Uncharacterized protein LOC126991007 [*Eriocheir sinensis*] | -4.373 | 8.66E-05 | 1.61E-02 |
| TRINITY_DN11402_c0_g2_i1 | Zinc finger protein [*Armadillidium nasatum*] | -4.383 | 8.32E-05 | 1.58E-02 |
| TRINITY_DN16_c1_g1_i3 | Putative ZDHHC-type Palmitoyltransferase 6 [*Armadillidium nasatum*] | -4.383 | 8.96E-05 | 1.64E-02 |
| TRINITY_DN1937_c0_g3_i11 | Hypothetical protein [*Idotea baltica*] | -4.411 | 4.82E-05 | 1.03E-02 |
| TRINITY_DN1073_c0_g1_i5 | Zinc finger protein 277-like [*Penaeus japonicus*] | -4.428 | 8.10E-05 | 1.55E-02 |
| TRINITY_DN1939_c1_g1_i20 | Serine/threonine-protein kinase D3-like isoform X1 [*Cherax quadricarinatus*] | -4.435 | 7.62E-05 | 1.49E-02 |
| TRINITY_DN1107_c0_g1_i5 | #N/A | -4.435 | 5.16E-05 | 1.08E-02 |
| TRINITY_DN1208_c0_g1_i10 | Hypothetical protein Avbf_09238 [*Armadillidium vulgare*] | -4.444 | 1.36E-04 | 2.27E-02 |
| TRINITY_DN1849_c0_g3_i6 | Putative fatty acyl-CoA reductase [*Armadillidium nasatum*] | -4.447 | 8.68E-05 | 1.61E-02 |
| TRINITY_DN1747_c0_g1_i13 | Galactose-1-phosphate uridylyltransferase [*Armadillidium nasatum*] | -4.451 | 6.36E-05 | 1.30E-02 |
| TRINITY_DN5724_c2_g1_i13 | NGFI-A-binding protein homolog isoform X16 [*Portunus trituberculatus*] | -4.484 | 1.93E-04 | 2.98E-02 |
| TRINITY_DN17173_c0_g1_i15 | Uncharacterized protein LOC119584321 [*Penaeus monodon*] | -4.487 | 1.57E-04 | 2.55E-02 |
| TRINITY_DN3845_c0_g1_i16 | ERI1 exoribonuclease 2-like [*Penaeus japonicus*] | -4.489 | 9.37E-05 | 1.71E-02 |
| TRINITY_DN1345_c0_g1_i10 | #N/A | -4.500 | 4.52E-05 | 1.02E-02 |
| TRINITY_DN216768_c0_g1_i1 | #N/A | -4.525 | 5.16E-05 | 1.08E-02 |
| TRINITY_DN576_c0_g1_i11 | Hypothetical protein Anas_07065 [*Armadillidium nasatum*] | -4.526 | 3.86E-05 | 8.85E-03 |
| TRINITY_DN56_c6_g3_i4 | #N/A | -4.530 | 4.95E-05 | 1.05E-02 |
| TRINITY_DN3086_c0_g1_i10 | Calcium-transporting ATPase [*Cherax cainii*] | -4.549 | 8.98E-05 | 1.64E-02 |
| TRINITY_DN594_c0_g1_i24 | #N/A | -4.575 | 4.72E-05 | 1.03E-02 |
| TRINITY_DN5147_c0_g1_i41 | #N/A | -4.595 | 2.92E-04 | 4.14E-02 |
| TRINITY_DN19403_c0_g2_i7 | Programmed cell death protein 4 [*Armadillidium nasatum*] | -4.598 | 8.98E-05 | 1.64E-02 |
| TRINITY_DN4665_c1_g1_i4 | Transcriptional enhancer factor TEF-1-like isoform X2 [*Penaeus monodon*] | -4.619 | 1.76E-04 | 2.78E-02 |
| TRINITY_DN2653_c0_g1_i35 | Ribonuclease 3 [*Armadillidium vulgare*] | -4.629 | 5.55E-05 | 1.15E-02 |
| TRINITY_DN874_c3_g1_i2 | #N/A | -4.645 | 4.35E-05 | 9.90E-03 |
| TRINITY_DN3627_c0_g2_i5 | Abnormal spindle-like microcephaly-associated protein homolog [*Penaeus vannamei*] | -4.672 | 2.40E-05 | 5.93E-03 |
| TRINITY_DN4792_c0_g1_i3 | Hypothetical protein [*Idotea baltica*] | -4.682 | 1.11E-04 | 1.93E-02 |
| TRINITY_DN3040_c0_g4_i2 | Thyroid receptor-interacting protein 6-like isoform X2 [*Portunus trituberculatus*] | -4.711 | 2.91E-05 | 6.94E-03 |
| TRINITY_DN895_c0_g1_i12 | Erythroid differentiation-related factor 1 [*Armadillidium nasatum*] | -4.729 | 1.96E-05 | 5.03E-03 |
| TRINITY_DN3286_c0_g2_i21 | Ras association domain-containing protein 5, partial [*Armadillidium nasatum*] | -4.733 | 8.70E-05 | 1.61E-02 |
| TRINITY_DN11748_c0_g2_i1 | #N/A | -4.771 | 1.44E-04 | 2.39E-02 |
| TRINITY_DN1603_c0_g1_i5 | Histone H4-like [*Daphnia pulicaria*] | -4.827 | 1.45E-05 | 4.02E-03 |
| TRINITY_DN439_c1_g1_i7 | Putative DNA helicase MCM8-like [*Penaeus vannamei*] | -4.849 | 1.47E-05 | 4.05E-03 |
| TRINITY_DN769_c0_g1_i24 | #N/A | -4.855 | 1.65E-05 | 4.39E-03 |
| TRINITY_DN3257_c0_g1_i1 | Hypothetical protein [*Idotea baltica*] | -4.886 | 1.42E-05 | 3.97E-03 |
| TRINITY_DN5226_c0_g2_i4 | Thioredoxin reductase 3 [*Armadillidium nasatum*] | -4.933 | 9.47E-06 | 2.98E-03 |
| TRINITY_DN3373_c0_g3_i5 | Putative RNA-directed DNA polymerase from transposon X-element [*Penaeus vannamei*] | -4.933 | 1.51E-04 | 2.47E-02 |
| TRINITY_DN51227_c0_g2_i1 | Ribokinase [*Armadillidium nasatum*] | -4.941 | 1.15E-04 | 1.99E-02 |
| TRINITY_DN1768_c1_g2_i1 | Heat shock protein 27 isoform X2 [*Hyalella azteca*] | -4.954 | 1.02E-05 | 3.08E-03 |
| TRINITY_DN2479_c0_g1_i5 | Zinc finger H2C2-type histone UAS binding [*Trinorchestia longiramus*] | -4.955 | 8.92E-06 | 2.90E-03 |
| TRINITY_DN4924_c0_g1_i1 | Hypothetical protein Anas_04384 [*Armadillidium nasatum*] | -4.963 | 9.32E-06 | 2.98E-03 |
| TRINITY_DN24918_c0_g1_i9 | N-terminal kinase-like protein [*Armadillidium nasatum*] | -4.993 | 9.62E-05 | 1.73E-02 |
| TRINITY_DN665_c0_g1_i6 | Uncharacterized protein APZ42_003968 [*Daphnia magna*] | -5.003 | 1.43E-05 | 3.98E-03 |
| TRINITY_DN3710_c0_g1_i10 | Serine proteinase stubble [*Armadillidium nasatum*] | -5.021 | 3.25E-05 | 7.65E-03 |
| TRINITY_DN8538_c0_g1_i3 | Ankyrin repeat and LEM domain-containing protein 1, partial [*Armadillidium nasatum*] | -5.111 | 1.47E-05 | 4.05E-03 |
| TRINITY_DN234_c0_g1_i1 | G2/mitotic-specific cyclin-A-like [*Penaeus chinensis*] | -5.145 | 2.34E-05 | 5.81E-03 |
| TRINITY_DN25964_c0_g1_i2 | Nucleoporin GLE1 [*Armadillidium nasatum*] | -5.194 | 1.20E-04 | 2.05E-02 |
| TRINITY_DN9955_c0_g1_i2 | Golgin subfamily A member 1 [*Armadillidium nasatum*] | -5.220 | 4.69E-05 | 1.03E-02 |
| TRINITY_DN3342_c2_g1_i6 | Cytochrome P450 9e2, partial [*Armadillidium vulgare*] | -5.237 | 2.85E-05 | 6.83E-03 |
| TRINITY_DN4538_c0_g1_i6 | Peroxisome proliferator-activated receptor gamma coactivator 1-alpha, partial [*Apis cerana*] | -5.268 | 1.23E-05 | 3.53E-03 |
| TRINITY_DN3551_c0_g1_i2 | D-3-phosphoglycerate dehydrogenase [*Armadillidium nasatum*] | -5.325 | 6.59E-06 | 2.23E-03 |
| TRINITY_DN1716_c2_g1_i3 | #N/A | -5.404 | 2.19E-06 | 9.13E-04 |
| TRINITY_DN15805_c0_g1_i3 | Uncharacterized protein LOC128684166 [*Cherax quadricarinatus*] | -5.443 | 9.42E-06 | 2.98E-03 |
| TRINITY_DN11611_c0_g2_i4 | Hypothetical protein [*Idotea baltica*] | -5.485 | 6.27E-06 | 2.16E-03 |
| TRINITY_DN4387_c1_g1_i3 | Uncharacterized protein LOC122252537 [*Penaeus japonicus*] | -5.494 | 8.50E-05 | 1.60E-02 |
| TRINITY_DN439_c1_g1_i1 | DNA helicase MCM8, partial [*Armadillidium nasatum*] | -5.502 | 2.45E-06 | 9.80E-04 |
| TRINITY_DN36286_c0_g1_i1 | Nucleoporin NUP53 [*Armadillidium vulgare*] | -5.517 | 2.25E-06 | 9.26E-04 |
| TRINITY_DN15702_c0_g1_i13 | Dynactin subunit 4-like [*Penaeus monodon*] | -5.530 | 2.79E-04 | 4.03E-02 |
| TRINITY_DN57159_c0_g1_i4 | Histone H2B [*Armadillidium nasatum*] | -5.570 | 1.31E-06 | 6.13E-04 |
| TRINITY_DN11193_c0_g1_i8 | Protein regulator of cytokinesis 1-like isoform X1 [*Penaeus chinensis*] | -5.579 | 2.38E-06 | 9.69E-04 |
| TRINITY_DN474_c0_g1_i10 | Uncharacterized protein LOC125029819 [*Penaeus chinensis*] | -5.603 | 2.96E-06 | 1.13E-03 |
| TRINITY_DN707_c2_g1_i21 | #N/A | -5.633 | 6.49E-06 | 2.21E-03 |
| TRINITY_DN4413_c1_g2_i1 | PDZ domain-containing protein GIPC1-like [*Penaeus monodon*] | -5.633 | 1.37E-06 | 6.33E-04 |
| TRINITY_DN5579_c0_g1_i5 | AFG3-like protein 2 [*Penaeus japonicus*] | -5.650 | 1.43E-06 | 6.49E-04 |
| TRINITY_DN911_c0_g1_i3 | Moesin/ezrin/radixin homolog 1-like isoform X3 [*Penaeus vannamei*] | -5.683 | 1.58E-06 | 6.90E-04 |
| TRINITY_DN1195_c0_g1_i1 | #N/A | -5.738 | 6.98E-07 | 3.46E-04 |
| TRINITY_DN7571_c0_g1_i1 | Diphthamide biosynthesis protein 2 [*Armadillidium nasatum*] | -5.794 | 3.44E-06 | 1.25E-03 |
| TRINITY_DN1092_c0_g1_i13 | Alpha-tocopherol transfer protein-like isoform X2 [*Homarus americanus*] | -5.866 | 4.70E-07 | 2.54E-04 |
| TRINITY_DN1664_c0_g1_i7 | #N/A | -5.923 | 1.46E-06 | 6.55E-04 |
| TRINITY_DN5579_c0_g2_i11 | AFG3-like protein 2 [*Procambarus clarkii*] | -6.007 | 5.34E-07 | 2.81E-04 |
| TRINITY_DN11586_c0_g1_i35 | Histone acetyltransferase KAT7, partial [*Armadillidium nasatum*] | -6.159 | 1.81E-05 | 4.73E-03 |
| TRINITY_DN31642_c0_g1_i1 | #N/A | -6.187 | 1.12E-05 | 3.28E-03 |
| TRINITY_DN8525_c0_g2_i3 | Iodotyrosine deiodinase-like isoform X3 [*Eriocheir sinensis*] | -6.340 | 1.45E-07 | 1.02E-04 |
| TRINITY_DN21794_c0_g1_i6 | N-acetyltransferase ESCO2 [*Armadillidium nasatum*] | -6.508 | 2.79E-04 | 4.03E-02 |
| TRINITY_DN723_c8_g2_i4 | #N/A | -6.627 | 1.96E-04 | 3.02E-02 |
| TRINITY_DN1811_c0_g2_i19 | #N/A | -6.671 | 1.50E-05 | 4.06E-03 |
| TRINITY_DN1363_c0_g1_i3 | Gastric triacylglycerol lipase-like [*Penaeus vannamei*] | -6.756 | 1.42E-04 | 2.35E-02 |
| TRINITY_DN8169_c0_g1_i14 | Protein-lysine N-methyltransferase n6amt2 [*Armadillidium nasatum*] | -6.778 | 1.28E-04 | 2.15E-02 |
| TRINITY_DN29088_c0_g2_i4 | Protein fem-1-like protein C [*Armadillidium nasatum*] | -6.852 | 1.04E-04 | 1.84E-02 |
| TRINITY_DN2316_c6_g1_i5 | #N/A | -6.865 | 1.04E-04 | 1.84E-02 |
| TRINITY_DN9833_c3_g2_i1 | #N/A | -6.913 | 8.59E-05 | 1.61E-02 |
| TRINITY_DN3113_c0_g1_i26 | Hypothetical protein [*Idotea baltica*] | -6.973 | 7.13E-05 | 1.42E-02 |
| TRINITY_DN9592_c0_g1_i4 | #N/A | -6.990 | 7.13E-05 | 1.42E-02 |
| TRINITY_DN13_c0_g1_i9 | Solute carrier family 12 member 6 [*Armadillidium nasatum*] | -7.021 | 6.51E-05 | 1.32E-02 |
| TRINITY_DN21006_c0_g1_i4 | Zinc finger protein [*Armadillidium vulgare*] | -7.092 | 5.01E-05 | 1.06E-02 |
| TRINITY_DN93012_c0_g1_i3 | Chitinase-3-like protein 2, partial [*Armadillidium nasatum*] | -7.102 | 5.34E-07 | 2.81E-04 |
| TRINITY_DN7997_c4_g1_i3 | #N/A | -7.113 | 5.01E-05 | 1.06E-02 |
| TRINITY_DN2262_c5_g1_i6 | Phosphatidylinositol-glycan biosynthesis class X protein [*Armadillidium vulgare*] | -7.127 | 4.60E-05 | 1.02E-02 |
| TRINITY_DN1839_c1_g1_i6 | Protein KIBRA [*Armadillidium vulgare*] | -7.141 | 4.60E-05 | 1.02E-02 |
| TRINITY_DN3373_c0_g3_i7 | Putative RNA-directed DNA polymerase from transposon X-element [*Penaeus vannamei*] | -7.171 | 4.24E-05 | 9.67E-03 |
| TRINITY_DN1603_c0_g1_i2 | Histone H4-like [*Aedes aegypti*] | -7.173 | 8.38E-09 | 9.56E-06 |
| TRINITY_DN11330_c0_g1_i1 | Neprilysin-2-like [*Penaeus monodon*] | -7.229 | 3.33E-05 | 7.75E-03 |
| TRINITY_DN10879_c0_g1_i2 | 5-Methylcytosine rRNA methyltransferase NSUN4-like isoform X1 [*Eriocheir sinensis*] | -7.252 | 3.33E-05 | 7.75E-03 |
| TRINITY_DN15670_c0_g2_i2 | Uncharacterized protein LOC119580297 [*Penaeus monodon*] | -7.305 | 2.65E-05 | 6.45E-03 |
| TRINITY_DN19618_c0_g1_i1 | Centrosomal protein [*Armadillidium vulgare*] | -7.315 | 2.65E-05 | 6.45E-03 |
| TRINITY_DN11527_c0_g1_i3 | Exosome complex component RRP43 [*Armadillidium nasatum*] | -7.324 | 7.04E-08 | 5.76E-05 |
| TRINITY_DN95273_c0_g1_i4 | Extended synaptotagmin-2-like isoform X3 [*Penaeus chinensis*] | -7.425 | 1.84E-05 | 4.75E-03 |
| TRINITY_DN10618_c1_g1_i7 | #N/A | -7.428 | 1.84E-05 | 4.75E-03 |
| TRINITY_DN19066_c0_g1_i5 | LOW-QUALITY PROTEIN: MAM and LDL-receptor class A domain-containing protein 1-like [*Penaeus vannamei*] | -7.442 | 1.84E-05 | 4.75E-03 |
| TRINITY_DN1327_c1_g1_i14 | Hypothetical protein [*Idotea baltica*] | -7.457 | 1.72E-05 | 4.51E-03 |
| TRINITY_DN6014_c3_g1_i1 | Uncharacterized protein LOC121878169 [*Homarus americanu*s] | -7.505 | 1.50E-05 | 4.06E-03 |
| TRINITY_DN24308_c0_g1_i10 | LOW-QUALITY PROTEIN: Uncharacterized protein LOC128688584 [*Cherax quadricarinatus*] | -7.509 | 1.41E-05 | 3.96E-03 |
| TRINITY_DN200_c12_g1_i2 | #N/A | -7.509 | 1.41E-05 | 3.96E-03 |
| TRINITY_DN7196_c0_g1_i3 | Pseudouridine-5'-phosphate glycosidase-like [*Penaeus chinensis*] | -7.528 | 1.41E-05 | 3.96E-03 |
| TRINITY_DN30122_c1_g1_i5 | UDP-N-acetylglucosamine transporter-like isoform X2 [*Penaeus japonicus*] | -7.579 | 1.16E-05 | 3.35E-03 |
| TRINITY_DN2539_c0_g1_i1 | Peroxidase skpo-1 [*Armadillidium vulgare*] | -7.584 | 1.16E-05 | 3.35E-03 |
| TRINITY_DN59490_c0_g1_i4 | DNA-directed primase/polymerase protein-like [*Cherax quadricarinatus*] | -7.592 | 1.09E-05 | 3.21E-03 |
| TRINITY_DN28099_c1_g1_i3 | #N/A | -7.605 | 9.62E-06 | 2.98E-03 |
| TRINITY_DN8261_c0_g3_i3 | LOW-QUALITY PROTEIN: piggyBac transposable element-derived protein 3-like [*Portunus trituberculatus*] | -7.608 | 1.09E-05 | 3.21E-03 |
| TRINITY_DN1182_c2_g1_i1 | #N/A | -7.611 | 1.02E-05 | 3.08E-03 |
| TRINITY_DN4769_c0_g1_i14 | Uncharacterized protein LOC119570384, partial [*Penaeus monodon*] | -7.618 | 1.02E-05 | 3.08E-03 |
| TRINITY_DN16091_c0_g1_i12 | Uncharacterized protein LOC123754561 [*Procambarus clarkii*] | -7.646 | 9.62E-06 | 2.98E-03 |
| TRINITY_DN74807_c0_g2_i4 | #N/A | -7.661 | 9.05E-06 | 2.93E-03 |
| TRINITY_DN30605_c0_g1_i15 | Mitogen-activated protein kinase kinase kinase kinase 5-like isoform X4 [*Penaeus chinensis*] | -7.679 | 8.53E-06 | 2.79E-03 |
| TRINITY_DN818_c0_g1_i4 | GTP-binding protein [*Armadillidium nasatum*] | -7.684 | 8.53E-06 | 2.79E-03 |
| TRINITY_DN47217_c0_g1_i39 | Synembryn-A-like [*Penaeus vannamei*] | -7.711 | 7.58E-06 | 2.52E-03 |
| TRINITY_DN69245_c0_g1_i1 | Hypothetical protein Avbf_06109 [*Armadillidium vulgare*] | -7.719 | 8.18E-10 | 1.50E-06 |
| TRINITY_DN1092_c0_g1_i23 | Retinol-binding protein pinta-like isoform X1 [*Portunus trituberculatus*] | -7.760 | 1.32E-09 | 2.04E-06 |
| TRINITY_DN5998_c0_g1_i16 | Hypothetical protein [*Idotea baltica*] | -7.764 | 6.39E-06 | 2.18E-03 |
| TRINITY_DN23949_c1_g1_i6 | #N/A | -7.784 | 6.04E-06 | 2.09E-03 |
| TRINITY_DN25847_c0_g1_i42 | #N/A | -7.840 | 5.13E-06 | 1.79E-03 |
| TRINITY_DN334_c2_g1_i1 | #N/A | -7.869 | 4.61E-06 | 1.64E-03 |
| TRINITY_DN22553_c0_g1_i10 | Gastrula zinc finger protein XlCGF8.2DB [*Armadillidium vulgare*] | -7.935 | 3.75E-06 | 1.35E-03 |
| TRINITY_DN3056_c0_g1_i1 | #N/A | -7.967 | 3.40E-06 | 1.25E-03 |
| TRINITY_DN12971_c0_g1_i28 | Armadillo-like helical domain-containing protein 3 [*Homarus americanus*] | -7.972 | 3.40E-06 | 1.25E-03 |
| TRINITY_DN4730_c2_g1_i17 | Rho guanine nucleotide exchange factor 26 [*Armadillidium nasatum*] | -7.977 | 3.23E-06 | 1.21E-03 |
| TRINITY_DN2988_c0_g1_i3 | Hypothetical protein [*Idotea baltica*] | -7.982 | 3.23E-06 | 1.21E-03 |
| TRINITY_DN626_c2_g3_i2 | #N/A | -7.997 | 3.08E-06 | 1.17E-03 |
| TRINITY_DN52229_c0_g1_i11 | Protein NRDE2-like protein, partial [*Armadillidium nasatum*] | -8.038 | 2.67E-06 | 1.05E-03 |
| TRINITY_DN4250_c0_g2_i2 | Paired amphipathic helix protein Sin3a [*Armadillidium vulgare*] | -8.066 | 1.15E-09 | 1.83E-06 |
| TRINITY_DN768_c1_g1_i1 | tRNA methyltransferase [*Armadillidium nasatum*] | -8.096 | 2.22E-06 | 9.20E-04 |
| TRINITY_DN5524_c0_g1_i9 | Protein AF-10 [*Armadillidium nasatum*] | -8.124 | 2.03E-06 | 8.53E-04 |
| TRINITY_DN2396_c0_g1_i8 | Target of rapamycin complex subunit lst8 [*Armadillidium nasatum*] | -8.220 | 1.50E-06 | 6.68E-04 |
| TRINITY_DN2032_c0_g2_i21 | Paxillin-like isoform X6 [*Portunus trituberculatus*] | -8.232 | 1.44E-06 | 6.50E-04 |
| TRINITY_DN20321_c0_g1_i14 | Protein FAM63A [*Armadillidium vulgare*] | -8.251 | 1.38E-06 | 6.33E-04 |
| TRINITY_DN15605_c0_g1_i7 | Zinc finger protein [*Armadillidium vulgare*] | -8.308 | 1.13E-06 | 5.34E-04 |
| TRINITY_DN11702_c0_g1_i1 | #N/A | -8.332 | 1.04E-06 | 4.97E-04 |
| TRINITY_DN48247_c0_g1_i57 | TATA element modulatory factor [*Armadillidium nasatum*] | -8.434 | 7.45E-07 | 3.66E-04 |
| TRINITY_DN11528_c0_g1_i2 | Hypothetical protein Anas_03507 [*Armadillidium nasatum*] | -8.477 | 6.69E-07 | 3.34E-04 |
| TRINITY_DN1171_c0_g1_i13 | #N/A | -8.479 | 6.45E-07 | 3.25E-04 |
| TRINITY_DN8622_c0_g1_i16 | Hypothetical protein [*Idotea baltica*] | -8.487 | 6.45E-07 | 3.25E-04 |
| TRINITY_DN5337_c0_g1_i19 | #N/A | -8.496 | 6.23E-07 | 3.19E-04 |
| TRINITY_DN29814_c0_g2_i1 | #N/A | -8.520 | 5.81E-07 | 3.03E-04 |
| TRINITY_DN57693_c0_g3_i6 | DNA endonuclease RBBP8 [*Armadillidium nasatum*] | -8.559 | 5.07E-07 | 2.72E-04 |
| TRINITY_DN9737_c0_g1_i31 | #N/A | -8.591 | 4.59E-07 | 2.50E-04 |
| TRINITY_DN64197_c0_g1_i15 | Leishmanolysin-like peptidase [*Procambarus clarkii*] | -8.628 | 4.03E-07 | 2.26E-04 |
| TRINITY_DN29886_c0_g1_i7 | #N/A | -8.660 | 3.67E-07 | 2.13E-04 |
| TRINITY_DN2907_c0_g1_i30 | Replication protein A 14 kDa subunit [*Armadillidium vulgare*] | -8.668 | 3.55E-07 | 2.09E-04 |
| TRINITY_DN774_c23_g1_i1 | Hypothetical protein NE865_04302 [*Phthorimaea operculella*] | -8.689 | 3.34E-07 | 1.98E-04 |
| TRINITY_DN874_c1_g1_i132 | Hypothetical protein Anas_00192 [*Armadillidium nasatum*] | -8.703 | 3.14E-07 | 1.88E-04 |
| TRINITY_DN21011_c1_g2_i2 | #N/A | -8.724 | 2.96E-07 | 1.79E-04 |
| TRINITY_DN505_c0_g1_i5 | #N/A | -8.727 | 2.96E-07 | 1.79E-04 |
| TRINITY_DN3289_c0_g2_i4 | #N/A | -8.778 | 2.48E-07 | 1.56E-04 |
| TRINITY_DN11054_c0_g1_i1 | Disks large-like protein 5 [*Armadillidium nasatum*] | -8.800 | 2.27E-07 | 1.45E-04 |
| TRINITY_DN11256_c1_g1_i8 | Hypothetical protein [*Idotea baltica*] | -8.846 | 1.97E-07 | 1.28E-04 |
| TRINITY_DN4004_c0_g2_i1 | Peroxidasin-like protein, partial [*Armadillidium nasatum*] | -8.852 | 1.92E-07 | 1.26E-04 |
| TRINITY_DN27372_c0_g1_i2 | #N/A | -8.878 | 1.77E-07 | 1.18E-04 |
| TRINITY_DN24393_c0_g1_i17 | Hypothetical protein Anas_02317 [*Armadillidium nasatum*] | -8.924 | 1.51E-07 | 1.04E-04 |
| TRINITY_DN13546_c0_g2_i13 | Inositol hexakisphosphate and diphosphoinositol-pentakisphosphate kinase 2-like isoform X5 [*Homarus americanus*] | -8.933 | 1.47E-07 | 1.02E-04 |
| TRINITY_DN3289_c0_g1_i13 | #N/A | -8.964 | 1.32E-07 | 9.42E-05 |
| TRINITY_DN1101_c1_g1_i19 | Protein-cysteine N-palmitoyltransferase HHAT-like [*Penaeus japonicus*] | -8.995 | 1.20E-07 | 8.83E-05 |
| TRINITY_DN707_c5_g1_i2 | Palmitoyl-protein thioesterase ABHD10, mitochondrial-like isoform X2 [*Homarus americanus*] | -9.003 | 1.17E-07 | 8.71E-05 |
| TRINITY_DN5965_c2_g1_i2 | #N/A | -9.097 | 8.52E-08 | 6.53E-05 |
| TRINITY_DN5953_c0_g3_i1 | #N/A | -9.108 | 8.33E-08 | 6.53E-05 |
| TRINITY_DN4373_c0_g1_i4 | Ras-related protein Rab-24 [*Armadillidium vulgare*] | -9.115 | 8.14E-08 | 6.53E-05 |
| TRINITY_DN5998_c0_g1_i10 | Hypothetical protein [*Idotea baltica*] | -9.186 | 6.35E-08 | 5.26E-05 |
| TRINITY_DN1899_c0_g1_i1 | 1 uncharacterized protein LOC126991007 [*Eriocheir sinensis*] | -9.191 | 6.35E-08 | 5.26E-05 |
| TRINITY_DN598_c0_g1_i5 | Peptide-N(4)-(N-acetyl-beta-glucosaminyl)asparagine amidase-like [*Penaeus chinensis*] | -9.223 | 5.69E-08 | 4.85E-05 |
| TRINITY_DN33509_c2_g3_i3 | #N/A | -9.226 | 5.57E-08 | 4.82E-05 |
| TRINITY_DN61920_c0_g1_i5 | SH3 domain-binding glutamic acid-rich-like protein [*Armadillidium nasatum*] | -9.266 | 4.91E-08 | 4.30E-05 |
| TRINITY_DN769_c0_g1_i12 | #N/A | -9.269 | 7.57E-10 | 1.43E-06 |
| TRINITY_DN26008_c0_g1_i4 | Poly(A) RNA polymerase gld-2 A-like [*Homarus americanus*] | -9.277 | 4.71E-08 | 4.19E-05 |
| TRINITY_DN2675_c0_g1_i4 | #N/A | -9.318 | 4.16E-08 | 3.76E-05 |
| TRINITY_DN160_c0_g1_i3 | Hypothetical protein [*Idotea baltica*] | -9.389 | 3.28E-08 | 3.01E-05 |
| TRINITY_DN474_c0_g1_i3 | Uncharacterized protein LOC125029819 [*Penaeus chinensis*] | -9.407 | 3.10E-08 | 2.93E-05 |
| TRINITY_DN2931_c0_g1_i9 | Multidrug resistance-associated protein 1-Like isoform X3 [*Penaeus japonicus*] | -9.445 | 2.72E-08 | 2.61E-05 |
| TRINITY_DN3627_c0_g2_i4 | abnormal spindle-like microcephaly-associated protein homolog [*Penaeus vannamei*] | -9.472 | 2.48E-08 | 2.42E-05 |
| TRINITY_DN4898_c0_g1_i1 | Protein cornichon-like [*Penaeus vannamei*] | -9.555 | 1.87E-08 | 1.95E-05 |
| TRINITY_DN4658_c0_g1_i10 | L-2-hydroxyglutarate dehydrogenase, mitochondrial [*Armadillidium nasatum*] | -9.662 | 1.32E-08 | 1.43E-05 |
| TRINITY_DN8813_c0_g1_i3 | Girdin [*Armadillidium vulgare*] | -9.684 | 1.22E-08 | 1.34E-05 |
| TRINITY_DN1278_c0_g1_i44 | Oocyte zinc finger protein XlCOF6.1 [*Armadillidium nasatum*] | -9.847 | 7.02E-09 | 8.67E-06 |
| TRINITY_DN1763_c0_g2_i1 | Hemolymph clottable protein [*Armadillidium nasatum*] | -9.905 | 7.58E-13 | 4.17E-09 |
| TRINITY_DN6125_c0_g1_i3 | #N/A | -9.939 | 5.20E-09 | 6.84E-06 |
| TRINITY_DN1252_c1_g1_i1 | Very low-density lipoprotein receptor, partial [*Armadillidium nasatum*] | -10.038 | 7.35E-13 | 4.17E-09 |
| TRINITY_DN7403_c1_g1_i6 | mpv17-like protein 2 [*Penaeus japonicus*] | -10.058 | 3.46E-09 | 4.76E-06 |
| TRINITY_DN6951_c0_g1_i21 | Kinesin-like protein KIF3B [*Armadillidium nasatum*] | -10.101 | 2.99E-09 | 4.21E-06 |
| TRINITY_DN12935_c0_g1_i5 | Carbohydrate-responsive element-binding protein-like isoform X2 [*Procambarus clarkii*] | -10.217 | 2.03E-09 | 2.93E-06 |
| TRINITY_DN293_c2_g1_i3 | Hypothetical protein Avbf_12335 [*Armadillidium vulgare*] | -10.404 | 1.09E-09 | 1.78E-06 |
| TRINITY_DN1416_c0_g1_i19 | Glutathione peroxidase-like isoform X1 [*Cherax quadricarinatus*] | -10.429 | 9.96E-10 | 1.67E-06 |
| TRINITY_DN3289_c0_g1_i2 | #N/A | -10.431 | 9.87E-10 | 1.67E-06 |
| TRINITY_DN8110_c0_g1_i6 | AT-rich interactive domain-containing protein 4B [*Penaeus vannamei*] | -10.463 | 8.90E-10 | 1.58E-06 |
| TRINITY_DN2134_c0_g1_i2 | Hypothetical protein Avbf_09547 [*Armadillidium vulgare*] | -10.588 | 5.16E-14 | 7.81E-10 |
| TRINITY_DN30125_c0_g1_i2 | Uncharacterized protein LOC119593038 [*Penaeus monodon*] | -10.609 | 5.43E-10 | 1.06E-06 |
| TRINITY_DN6210_c0_g1_i1 | Hypothetical protein Avbf_15556 [*Armadillidium vulgare*] | -10.650 | 4.72E-10 | 9.85E-07 |
| TRINITY_DN5618_c0_g2_i9 | Uncharacterized protein LOC122249196 isoform X4 [*Penaeus japonicus*] | -10.708 | 3.87E-10 | 8.62E-07 |
| TRINITY_DN1530_c0_g2_i2 | #N/A | -10.745 | 3.42E-10 | 7.95E-07 |
| TRINITY_DN145_c0_g1_i3 | #N/A | -10.801 | 1.67E-10 | 4.04E-07 |
| TRINITY_DN2314_c0_g1_i5 | #N/A | -11.000 | 1.45E-10 | 3.65E-07 |
| TRINITY_DN2483_c4_g1_i1 | Hypothetical protein Avbf_14903 [*Armadillidium vulgare*] | -11.274 | 1.07E-14 | 3.22E-10 |
| TRINITY_DN1252_c1_g2_i3 | #N/A | -11.304 | 5.15E-11 | 1.42E-07 |
| TRINITY_DN955_c0_g1_i17 | Hypothetical protein Anas_09068 [*Armadillidium nasatum*] | -11.403 | 3.68E-11 | 1.17E-07 |
| TRINITY_DN4139_c0_g1_i2 | Cytosolic phospholipase A2 [*Armadillidium vulgare*] | -11.433 | 3.33E-11 | 1.12E-07 |
| TRINITY_DN1605_c0_g1_i5 | Targeting protein for Xklp2 [*Armadillidium vulgare*] | -11.462 | 2.29E-11 | 8.16E-08 |
| TRINITY_DN3065_c1_g1_i5 | Period [*Eurydice pulchra*] | -12.338 | 1.53E-12 | 7.70E-09 |
| TRINITY_DN92127_c0_g2_i7 | Hypothetical protein Anas_13046 [*Armadillidium nasatum*] | -12.823 | 2.93E-13 | 1.97E-09 |
| TRINITY_DN154_c0_g1_i2 | U4/U6.U5 tri-snRNP-associated protein 1-like [*Cherax quadricarinatus*] | -12.951 | 1.89E-13 | 1.43E-09 |
| TRINITY_DN144_c2_g5_i1 | #N/A | -13.166 | 9.09E-14 | 1.10E-09 |
| TRINITY_DN1659_c0_g1_i11 | LOW-QUALITY PROTEIN: beta-galactosidase-1-like protein 2 [*Portunus trituberculatus*] | -13.598 | 2.08E-14 | 4.20E-10 |
| TRINITY_DN92127_c0_g2_i3 | Hypothetical protein Anas_13046 [*Armadillidium nasatum*] | -14.657 | 5.60E-16 | 3.39E-11 |

**Table S2. Relative abundance of the biomes classified by CA.**

Most biomes were described (p ≤ 0.1). Abbreviations: Control, group without microplastic (EPS) treatment; EPS, group without MP treatment; C_AV, average of the control group; EPS_AV, average of the EPS group; F value, value calculated by the F test; and P value, p value calculated by the t-test or Mann–Whitney U test, which was selected based on the results of the Shapiro–Wilk test and F test.

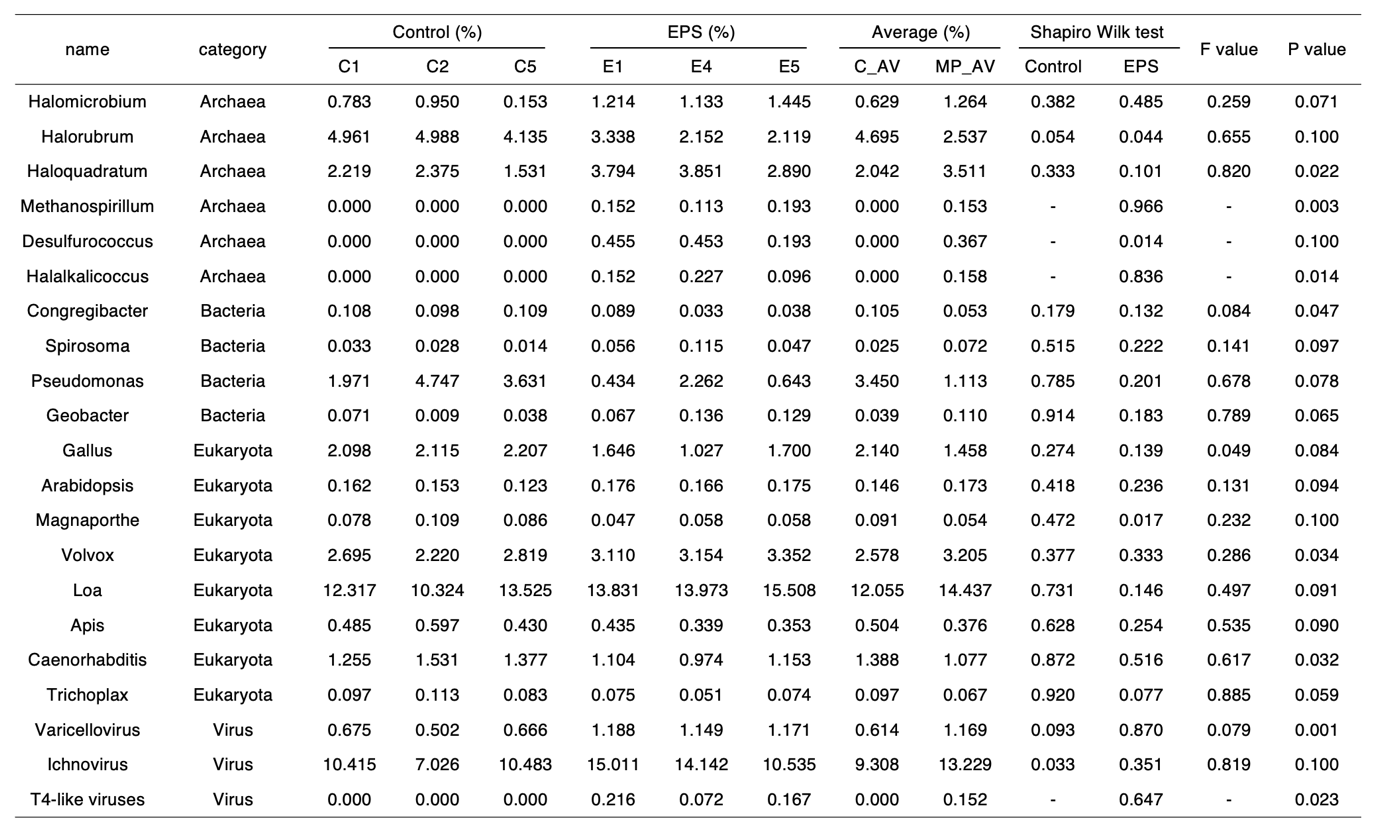

**Table S3. Indices calculated using EFA.**

EFA was calculated based on Minres (minimum residual). The other values are indicated as follows: EPS, group without MP treatment; h2, communality score; u2, uniqueness score; com, complexity, information score that is generally related to uniqueness.

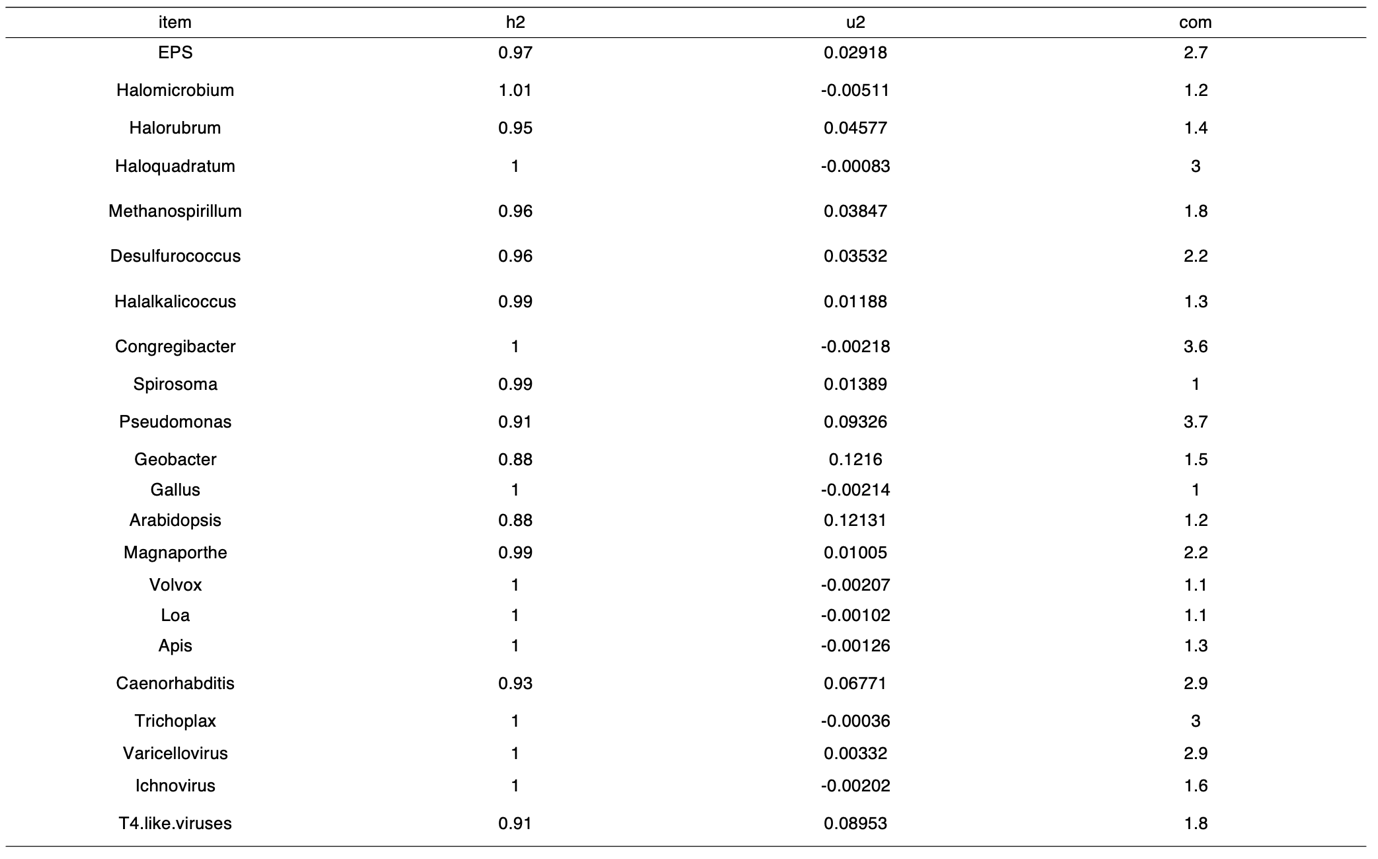

**Table S4. Statistical values of the final optimal structural equation models for symbiotic bacteria in wharf roach with EPS treatment.**

The bacteria used as the SEM components were selected from CA, RF, and EFA. Column no. 1 shows the best numerical SEM in Fig. 6b. Columns 2–4 show the models of inferior numerical structural equations. The inferior indices are underlined. Abbreviations: Test, EPS treatment; chisq, chi-square χ^2^; df, degrees of freedom; p value, p values (chi-square); cfi, comparative fix index (CFI); tli, Tucker–Lewis index (TLI); nfi, normed fit index (NFI); rfi, relative fit index (RFI); srmr, standardized root mean residuals (SRMR); AIC, Akaike information criterion; rmsea, root mean square error of approximation (RMSEA); gfi, goodness-of-fit index (GFI); and agfi, adjusted goodness-of-fit index (AGFI). The underlines are shown as values inferior to those of the other models with goodness-of-fit indices.

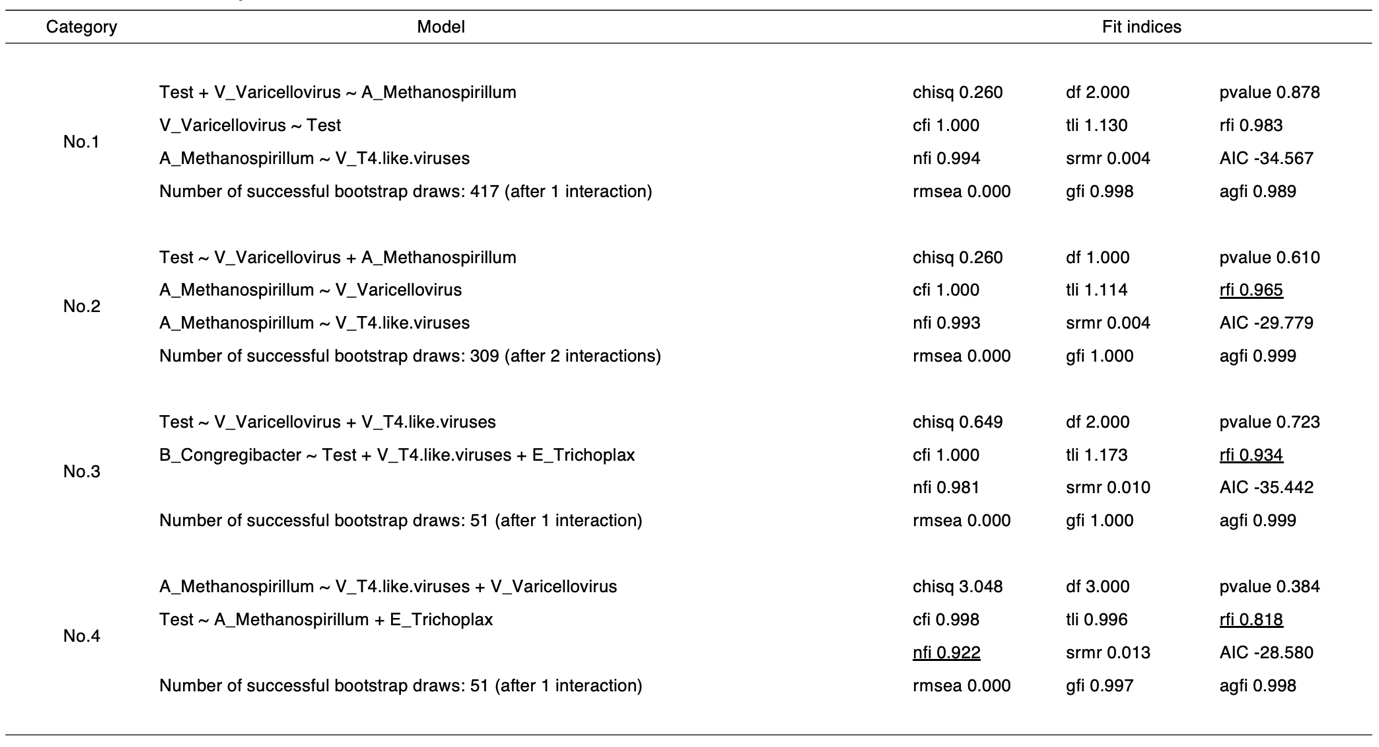

**Table S5.** List of models targeted by causal mediation analysis for Fig. 6b and their statistical values. The coefficients and calculated data related to the regression model are shown on the left-hand side of the figure. The values mediated by the regression models are shown on the right side. The number in “Stimulations” indicates the bootstrapping frequency. *, p < 0.05; **, p < 0.01; ***, p < 0.001.

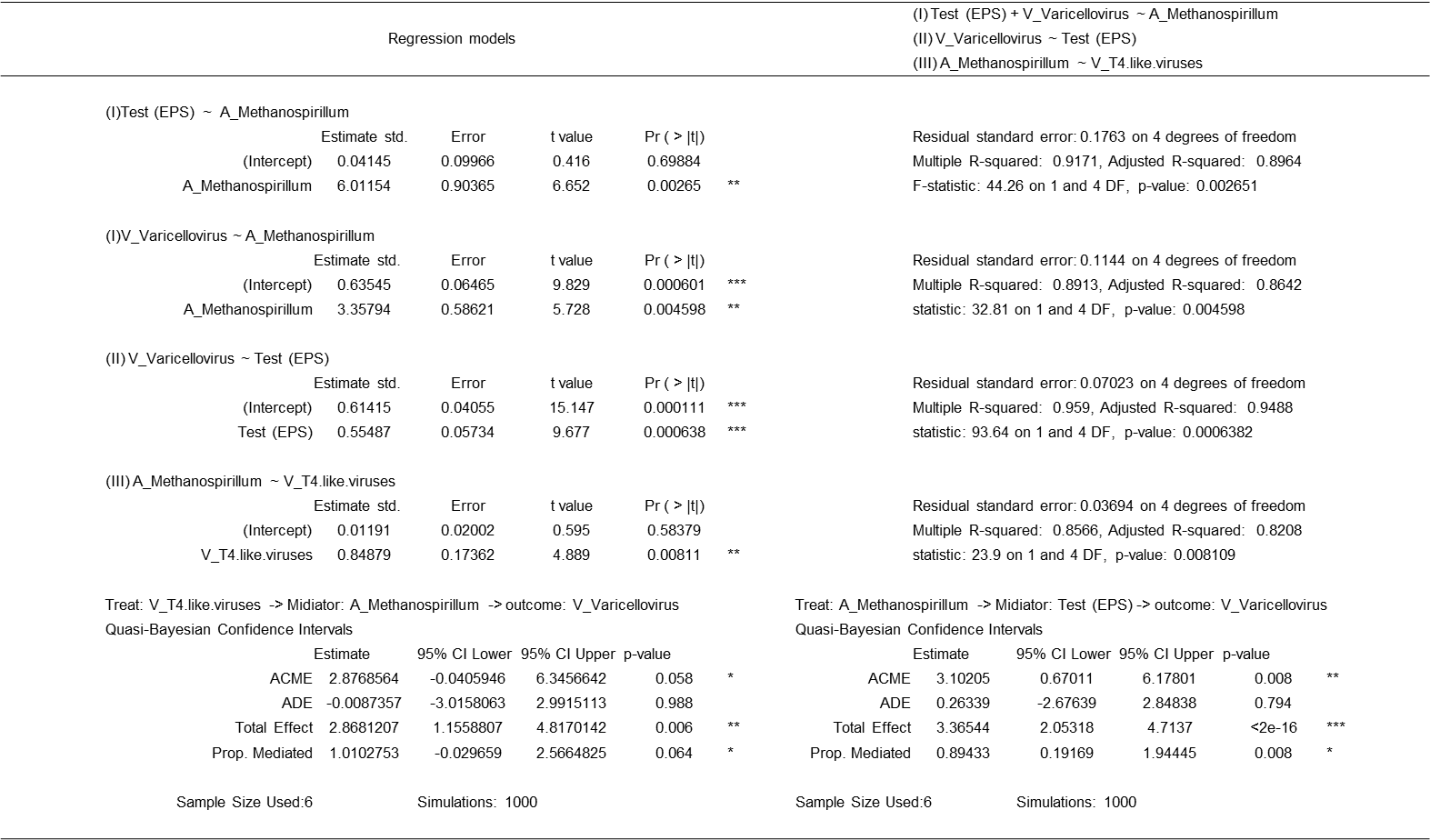

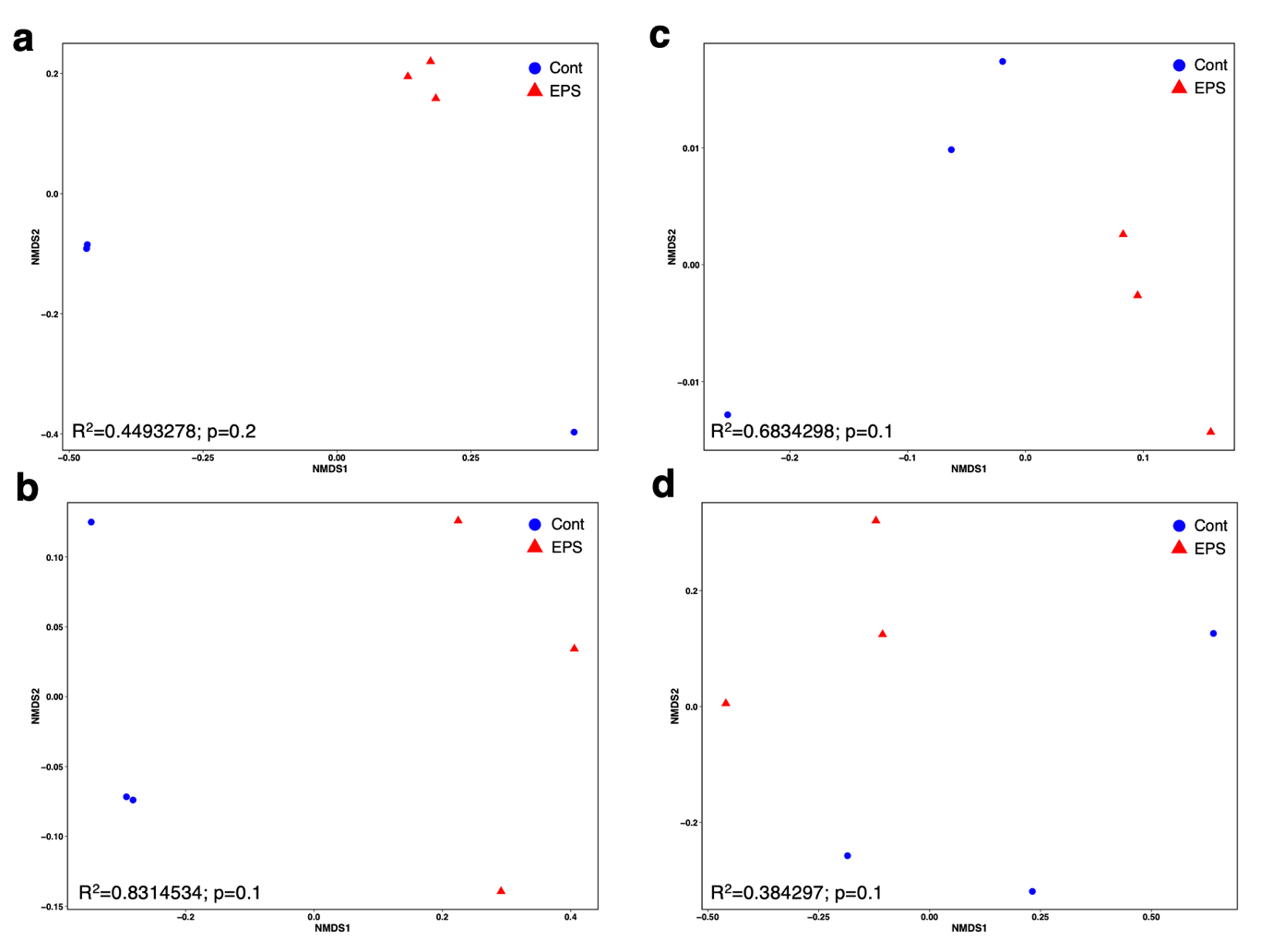

**Fig. S1. Non-metric multidimensional scaling (NMDS) plots of the biomes from the wharf roach.**

NMDS values were calculated using Bray–Curtis dissimilarity of the R library “vegarn.” R^2^ and p values were calculated using the R library “pairwiseAdonis.” The plots based on the data from (a) bacteria, (b) archaea, (c) eukaryota, and (d) viruses were visualized by the R library “ggplot2.”

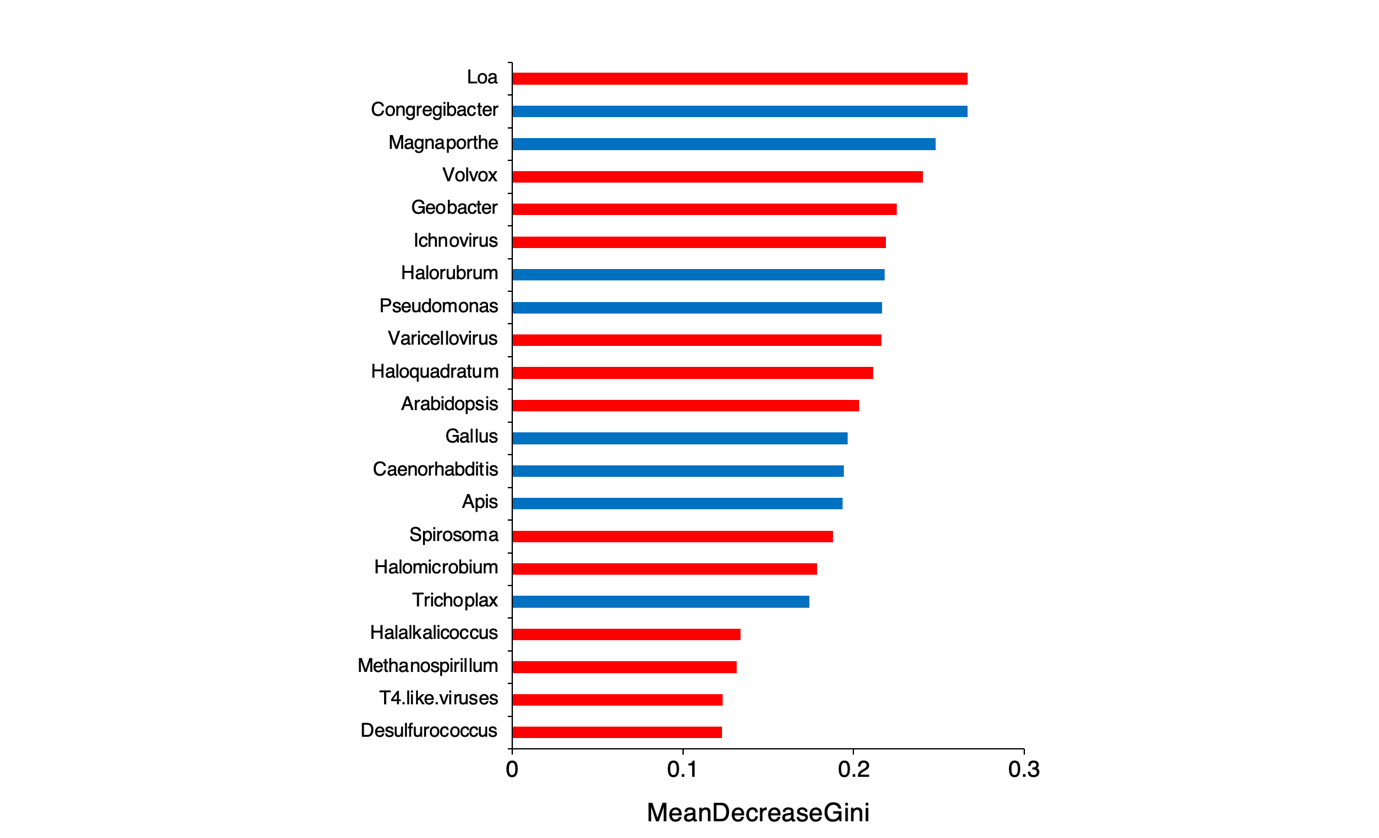

**Fig. S2. Extraction of feature factors by ML.**

Feature factors of the symbiotic biomes of *Ligia* spp. were selected using random forest. “MeanDecreasGini” means the value of feature factors. The blue bars show the components that increased under conditions without EPS treatment, and the red bars show the components that increased under conditions with EPS treatment.

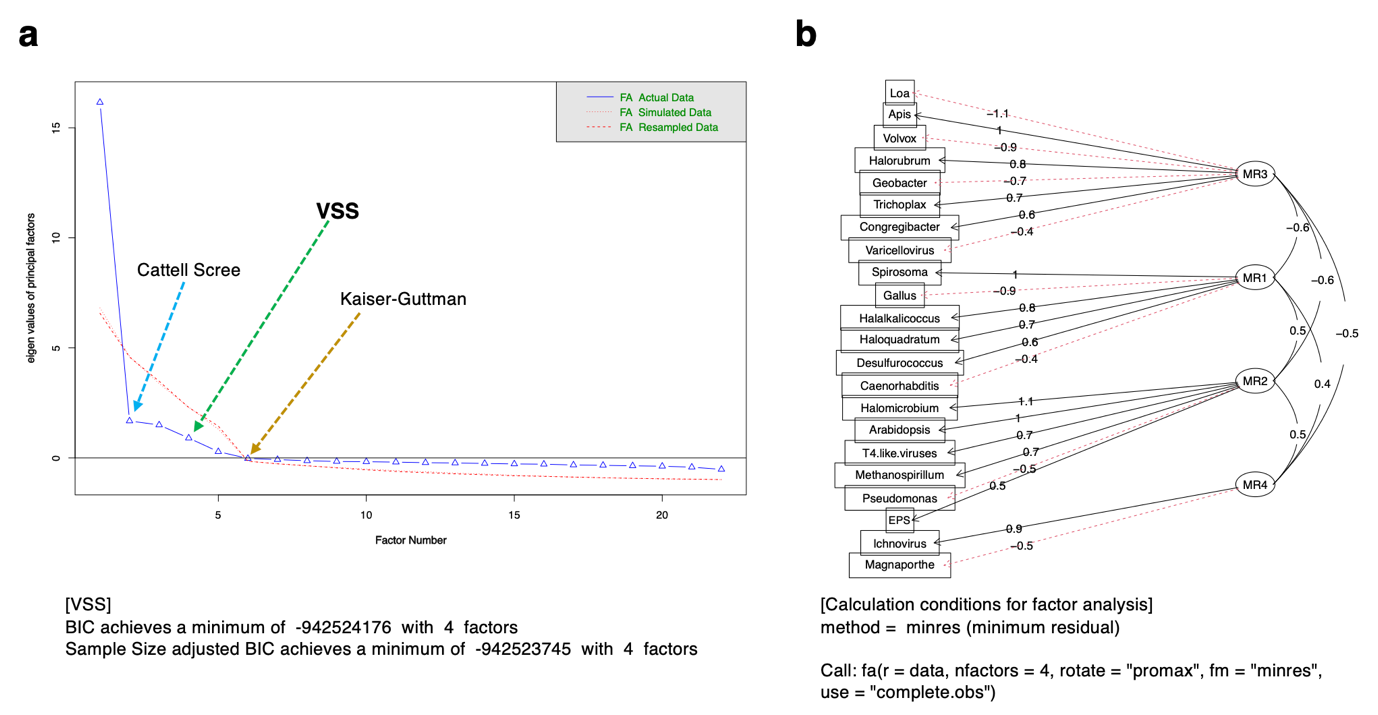

**Fig. S3. Evaluation by exploratory factor analysis (EFA) of the feature components.**

Panel (a) shows the parallel analysis scree plot for the EFA. The arrow point marked with “Catell Scree” (blue broken line) indicates the number of components based on the Cattle Scree test. The arrow point marked with “Kaiser–Guttman” (brown line) indicates the number of components based on the Kaiser–Guttman criteria. The arrow marked with “VSS” (green line) indicates the number of components based on the VSS-calculated criteria. (b) Components obtained from EFA setting 4 as factor number were visualized based on the upper calculation conditions. Abbreviations: FA, factor analysis; VSS, very simple structure; BIC, Bayesian information criterion; MR, minres (minimum residual) method; fa (r = data, nfactors = 4, rotate = “promax,” fm = “minres,” and use = “complete.obs”), calculation conditions for EFA.

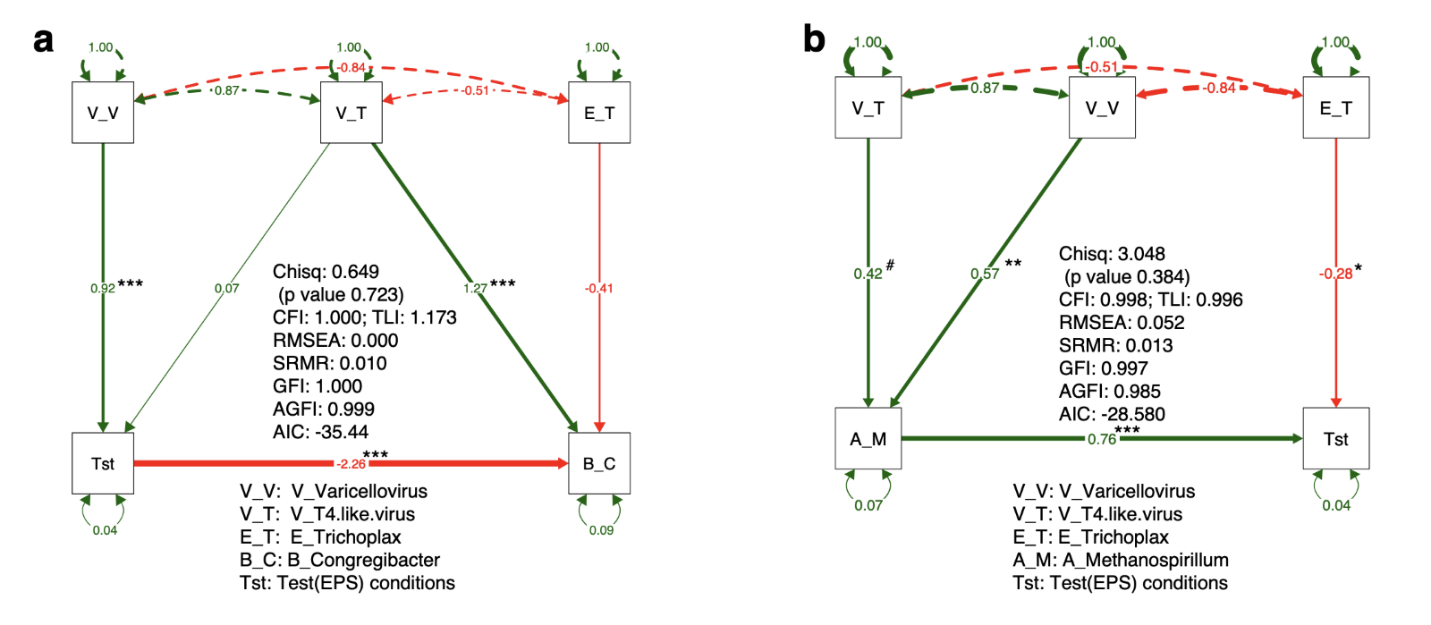

**Fig. S4. Visualization of the inferior component groups using structural equations.**

The two types of bacteria and eukaryotes were estimated using structural equation modeling (SEM) for confirmatory factor analysis (CFA). Models (a) No. 3 and (b) No. 4 in Table S6 were visualized by the R library “semPlot.” The color is indicated as follows: green, putative positive interaction; red, putative negative interaction. Abbreviations: Tst, conditions of the EPS administration; A_M, A_Methanospirillum; B_C, B_Congregibacter; E_T, E_Trichoplax; V_T, V_T4.like.virus; V_V, V_Varicellovirus. The main fit indices of the models are shown in the figures, and the other indices are shown in Table S2. Abbreviations of the main fit indices are as follows: chisq, Chi-square: χ^2^; p value, p values (Chi-square); CFI, comparative fix index; TLI, Tucker–Lewis index; RMSEA, root mean square error of approximation; SRMR, standardized root mean residuals; GFI, goodness-of-fit index; and AGFI, adjusted goodness-of-fit index. ^#^, p < 0.1; *, p < 0.05; **, p < 0.01; ***, p < 0.001.

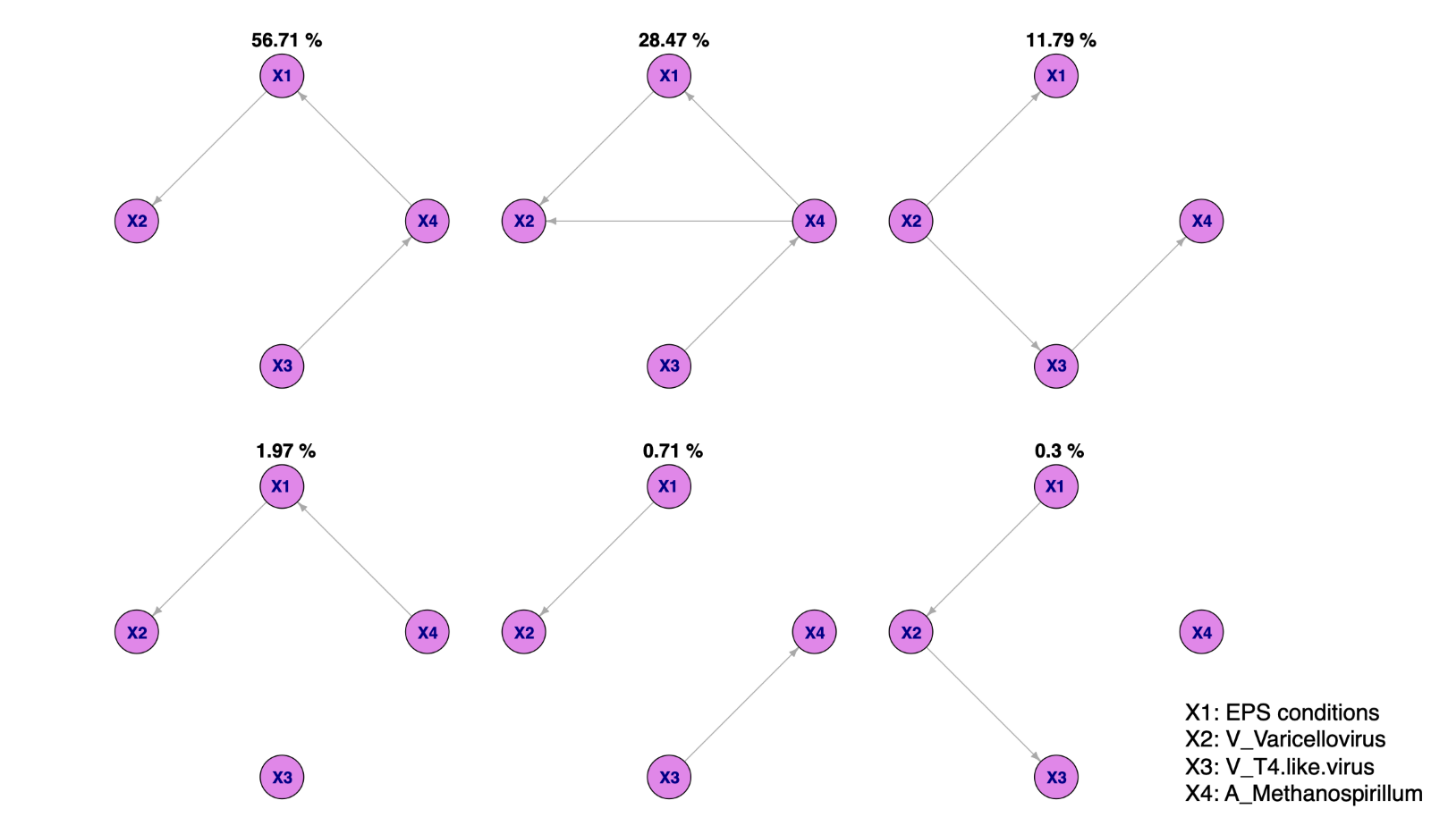
 **Fig. S5. Causal interaction networks of SEM-selected components predicted by BayesLiNGAM**.

The number at the top of each DAG represents the calculated percentage of the frequency of the DAG pattern. Abbreviations: X1, conditions with EPS treatment; X2, V_Varicellovirus; X3, V_T4.like.virus; X4, A_Methanospirillum.
